# A Unified Theory and Bayesian Framework for Phenological Inference from Biocollection Data: Resolving Paradoxes by Considering Phenophase Duration

**DOI:** 10.1101/2025.06.20.660820

**Authors:** David J. Hearn, Daniel S. Caetano

## Abstract

- Phenology, the study of recurring biological events, has gained increased attention with the rise of digitized biocollections. However, no general theoretical framework has linked variation in biocollection data to underlying phenological processes.
- We present a unified statistical theory of phenological distributions that mathematically integrates phenological extremes, onset timing, phenophase duration, cessation, and peak activity and links these phenomena to variation in biocollection data.
- A key insight from this theory is that phenological sensitivity estimates from biocollection data are confounded by phenophase duration. Both onset and duration contribute to variation in collection dates, and standard regression methods cannot disentangle these effects.
- To address this, we develop a Bayesian inference framework based on Gaussian processes (GPs) that explicitly models onset and duration as latent variables. We assess its performance under various scenarios, including model misspecification and identifiability, and show that it outperforms three alternative methods.
- Applying the method to an empirical dataset of over 5,000 herbarium records, we find that phenophase duration frequently acts as a confounder, correlating with one or more covariates across all species examined.
- These results highlight that phenophase duration is a critical but underappreciated factor in phenological studies using biocollection data.

## Introduction

Phenology is the study of biological processes that recur periodically. Leafing-out, flowering, fruiting, dormancy (or activity), and migration name a few phenological processes. Its study is important to understand the coordination of ecosystem processes mediated by biological activities (Fitter & Fitter, 2002; Sherry *et al*., 2007; Thackeray *et al*., 2016a; Renner & Zohner, 2018; Zohner *et al*., 2018). Phenology influences flows within biogeochemical cycles and impacts resource availability, more broadly. Within the past few decades, the digitization of biocollections together with national networks that track phenology (Crimmins, 2021) have made phenological data available at an unprecedented global spatial scale and temporal scale of multiple centuries. In particular, the digitization of plant specimens preserved in herbaria provides timing (dates of collection), locality of collection (latitude and longitude), and phenological state (based on analyses of digitized images of the physical specimens themselves) over several centuries of global collection. This digitization effort has made it possible to research broad-scale questions that would not be possible otherwise (Williams *et al*., 2021; Ahlstrand *et al*., 2025).

No general theory of phenological timing is available to model and analyze phenological data from biocollections. Our goal is to remedy this situation by providing a unified theory that connects phenological patterns to information gleaned from specimens. We focus specifically on the statistical and theoretical underpinnings needed to translate information from biocollection data into phenological insights.

Interest in the use of museum specimens to document phenological change and explore evolutionary and ecological ramifications extends across multiple decades. The following is a snapshot of this literature, focused on studies that used data from herbarium specimens: (Primack *et al*., 2004a; Miller-Rushing *et al*., 2006; Robbirt *et al*., 2011; Panchen *et al*., 2012; Lavoie, 2013; Calinger *et al*., 2013; Davis *et al*., 2015; Beauvais *et al*., 2017; Willis *et al*., 2017; Meineke & Davies, 2019; Park *et al*., 2019; Heberling, 2022; Iwanycki Ahlstrand *et al*., 2022; Bates *et al*., 2023; Park *et al*., 2025; Ahlstrand *et al*., 2025). One theme has been the expansion of datasets to investigate broadscale patterns at continental or global levels (Ahlstrand *et al*., 2025), even using over one million specimens to do so (Ramirez-Parada *et al*., 2024).

Coupled with this interest in the use of biocollection data is a growing literature on methods to use museum specimens for phenological research (Lavoie & Lachance, 2006; Moussus *et al*., 2010; Fitchett *et al*., 2015; Davis *et al*., 2015; Pearse *et al*., 2017; Jones & Daehler, 2018; Pearson, 2019a; Taylor, 2019; Belitz *et al*., 2020; Iler *et al*., 2021; Park *et al*., 2023; Wilson *et al*., 2023; Park *et al*., 2025; Lai, 2025). Many of these methods use regression techniques to fit models to observed dates of collection, e.g: (Primack *et al*., 2004b; Calinger *et al*., 2013; Davis *et al*., 2015). Earlier studies employed standard linear regression (SR) techniques directly to the day of year (DOY) dates extracted from herbarium specimens. Such regression models incorporated covariates, such as climate and spatial variables, to tease apart factors that contribute to variation in timing of phenological events, such as flowering and leafing out. More recent models add additional sophistication (McElreath, 2020; Willems *et al*., 2022; Bates *et al*., 2023; Park *et al*., 2025), but they still rely on fitting the model directly to the observed data. In particular, Willems et al. (Willems *et al*., 2022) recognize how inclusion of spatial covariates can influence inferences about regression slopes.

One of the key results of previous analyses has been the estimation of phenological sensitivities (Calinger *et al*., 2013; Wang *et al*., 2015; Thackeray *et al*., 2016a,b; Park *et al*., 2019; Meineke *et al*., 2021; Mazer *et al*., 2021; Xie *et al*., 2022). Phenological sensitivities describe how phenological processes change in tandem with climate or other covariates, and they quantify the strength of the impact of, e.g., climate change on phenological change. Many studies thus focused on how climate variables influence the onset of a phenological process (Abu-Asab *et al*., 2001; Primack *et al*., 2004b; Park & Mazer, 2018; Willems *et al*., 2022).

Phenological sensitivities are the estimated slope coefficients of covariates in the models. Davis et al. (Davis *et al*., 2015) investigated whether slopes obtained from data based on herbarium specimens are comparable to those obtained from field observations. Their analyses suggested that the slopes were statistically no different between the datasets. Based, in part, on this result, they concluded that herbarium data provide reliable records of phenological change. Although data from biocollections may correlate with phenological state (Primack *et al*., 2004a; Robbirt *et al*., 2011; Calinger *et al*., 2013; Davis *et al*., 2015; Park *et al*., 2019; Pearson, 2019b; Heberling, 2022), it is not directly evident that conventional approaches to analyze data from biocollections result in accurate inferences of phenological parameters of interest in a consistent fashion. In particular, our results demonstrate that slope estimates from regression models using biocollection data are not directly comparable to those derived from field-based observations of onset due to differences in what the data represent and how phenophase duration confounds observed patterns. Slopes of models through collection dates are therefore not expected to match slopes of onset models unless durations remain constant on average.

To demonstrate this result, we build a general theory of phenology that mathematically connects distributions of key phenological events, including phenological extreme events, onset time of phenophases, durations of phenophases, cessation times of phenophases, peak phenophase, and population level total temporal span of phenophases. Using this framework, we show that both variation in onset timing and variation in duration contribute to observed variation in collection dates. When the influence of duration exceeds that of onset, the resulting signal in collection dates can paradoxically oppose the true underlying onset signal.

One of the key innovations of our model is the treatment of phenological events as latent variables, quantities that are not directly observed, yet influence observed outcomes. It has long been recognized that herbarium specimens are not typically collected with phenology in mind (Davis *et al*., 2015). As such, the timing of phenological events is not directly visible from herbarium records. Phenological timing is therefore hidden or latent. Consequently, the precise timing of phenological events is not explicitly recorded in herbarium data, rendering these events effectively hidden. Our theory introduces a rigorously derived probabilistic framework that links observed specimen collection times to the unobserved, or latent, timing of phenological events. From this framework, we develop a Bayesian inference approach implemented in Stan (Carpenter *et al*., 2017) via R (R Core Team, 2025) wrappers provided by the *cmdstanr* package (Gabry *et al*., 2025).

The model we develop is also hierarchical (e.g., Wilson *et al*., 2023), explicitly recognizing multiple scales of organization, from individuals to populations, that contribute to phenological variation. A key implication of this approach is that methods which are not population-size aware may fail to accurately estimate phenological extremes from biocollection data, as these extremes are inherently defined at the population level in our model.

Because our framework is novel and departs in key ways from previous assumptions— particularly regarding slope interpretation and model structure—we devote a substantial portion of this manuscript to validating the approach and comparing it to existing methods. Specifically, we focus on comparisons with SR, quantile regression (QR; Park *et al*., 2025), and Pearse et al.’s (Pearse *et al*., 2017) estimator (PE) for phenological extremes. To evaluate performance, we rely heavily on simulated datasets with known parameters, enabling a direct assessment of each method’s accuracy.

To further validate our methods, we examine model identifiability, inference under model misspecification, and the influence of prior distributions on inference accuracy. We conclude with an analysis of empirical data from 13 species of spring ephemeral wildflowers in the northeastern United States. We chose these species because of extensive interest in spring ephemerals and because the impacts of climate on phenology for such groups is well documented (Oldham, 1990; Abu-Asab *et al*., 2001; Badeck *et al*., 2004; Primack *et al*., 2004a; Calinger *et al*., 2013; Pearson, 2019c; Petrauski *et al*., 2019; Alecrim *et al*., 2023; Miller *et al*., 2023; Faidiga *et al*., 2023; Watson & Vuorisalo, 2024; Miller & Stuble, 2024; Leoschke). The primary goal of this analysis is to assess the extent to which phenophase duration influences regression model slopes in real-world biocollection data, and we expect these species to be useful indicators of the correspondence between phenology and climate.

### Conceptual framework

The theory derived here applies to any biological process – not just phenology in the traditional sense – with the following eight Properties (Many illustrated by Fig. **1**):

1. The phenological process (i.e. phenophase) occurs at the level of an individual that is part of a larger population.
2. The phenophase has a start point (onset, *O*) and a finite duration (*D*) for each individual. Therefore, the phenophase also has a finite end time (cessation, *C*) that is the sum of the onset time and duration.
3. It is possible to observe (i.e., measure) when an individual starts, is in, and finishes a phenophase.
4. The onset and cessation of the process occur within a broader, but still finite, time period that repeats. Phenology can be analyzed at different scales of organization. Pertaining to temporal scales, the phenophase may occur within a time period of a day (Edery, 2000), within monthly time periods (Raible *et al*., 2017), within years (e.g., temperate zone flowering or fruiting; Helm *et al*., 2013), or during some other repeating time period. Throughout the paper, this repeated interval of time will be referred to as the “time period” (TP) to distinguish it from other intervals of time, such as days within yearly TPs.
5. Onset, duration, and cessation times vary among individuals within the population. The causes of variation may be due to individual plastic responses to random or deterministic variation in the external environmental, due to random noise during an individual’s development, or due to evolutionary processes. Because of the uncertainty imposed by such random factors, we employ a probabilistic framework to model phenology.
6. The distributions of phenological events have reached probabilistic stationarity: while individual phenological events (onset, duration, cessation) may vary from one time period (TP) to the next, the population-level distributions of these events remain stable over time, unless altered by environmental or genetic changes. In terms of stochastic processes, this reflects statistical equilibrium, where the flow of individuals into and out of phenophase states is balanced over time. As such, any observed changes in phenological distributions are not due to the population drifting randomly towards an equilibrium. Instead, any changes in distribution are due to external drivers, such as environmental variation or evolutionary change.
7. The timing of each individual’s phenophase is statistically independent of the other individuals in the population.
8. When carrying out inferences about population-level phenological processes, observed times (e.g., collection dates) of an individual in a phenophase are a random sample of time points within phenophases. Properties 6 through 8 indicate that individuals are independent and identically distributed (IID assumptions). These are standard assumptions of many statistical procedures such as ANOVA.

**Fig. 1.**
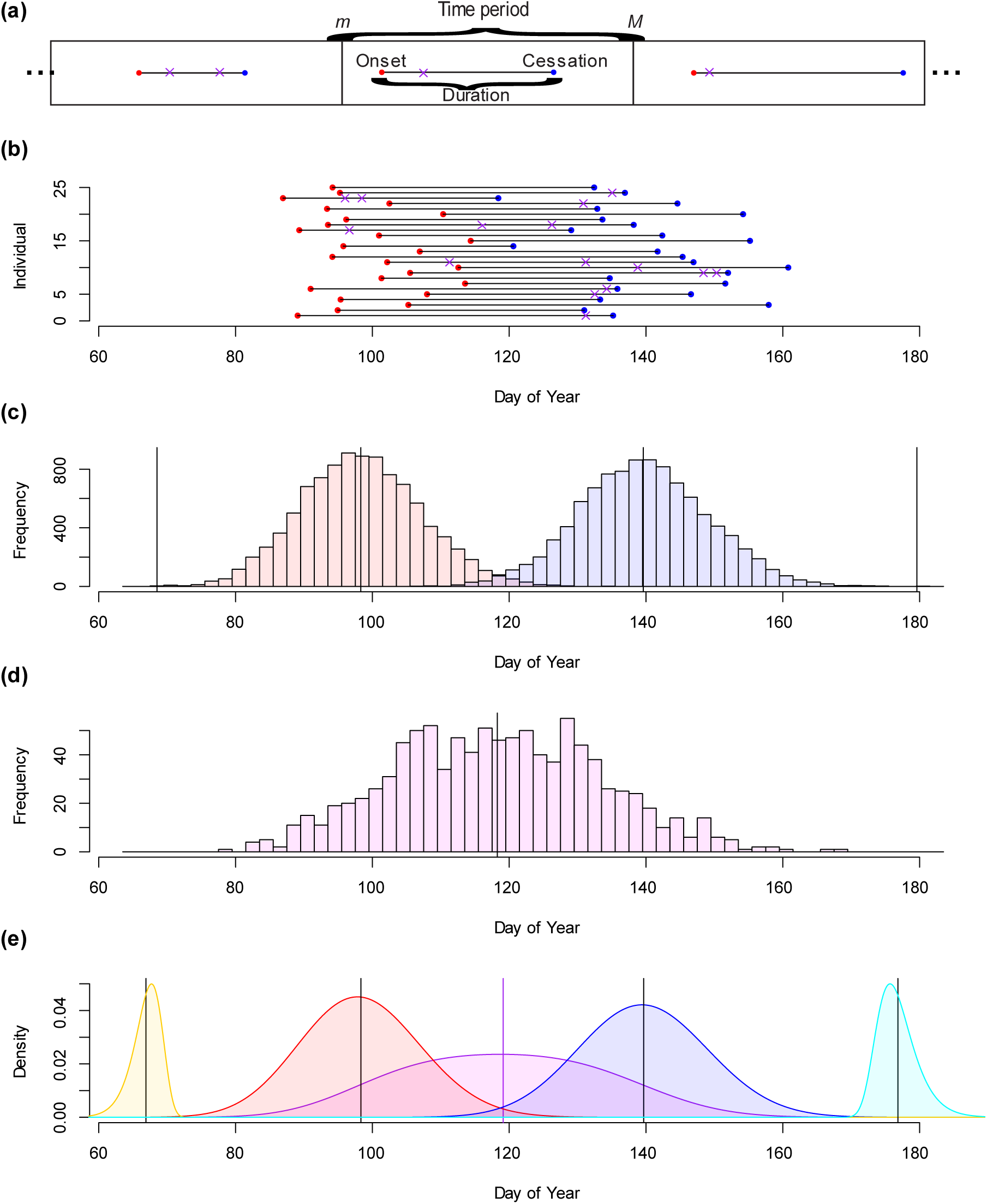
Conceptual framework for analyzing phenological phenomena at the population level. The color meaning is the same throughout all figures in this paper: red = onset, blue = cessation, gray = duration, purple = observed collection times. (a) Three repeating time periods (TPs) with one individual’s phenophase occurring in each period. ‘X’s represent observed collection times and the length of the line segment connecting the onset and cessation events is the phenophase duration. (b) A sample of 25 individuals from a population during one TP. Each individual may have a different onset, different duration, and different cessation. (c) Histograms of the unobserved (latent) onsets and cessations of all individuals in the population. Black vertical lines from left to right indicate the earliest onset, mean onset, mean cessation, and last cessation in the population. (d) The histogram of observed collection times (response variable *T*) of individuals in the phenophase. In biocollections, these are often the day of year (DOY) of collection of the specimens. The vertical line is the mean observed time. (e) Inferred probability density functions (PDFs) for the earliest onset (yellow), the onset (red), distribution of observed times (purple), cessation times (blue), and last cessation (cyan) inferred from the observed times illustrated in subfigure (d). Data for subfigures (b), (c), and (d) were simulated using procedures described in the text, as are inference procedures to obtain densities in (e).

### Formalization

Because there is variation within individuals across TPs, variation among individuals within a population, and intrinsic random noise that influences developmental outcomes, (Properties 5 and 6), we treat phenological variables (e.g., *O*, *D*, *C*) as random variables (RVs). Table 1 provides a full listing of notation and definitions.

**Table 1.**
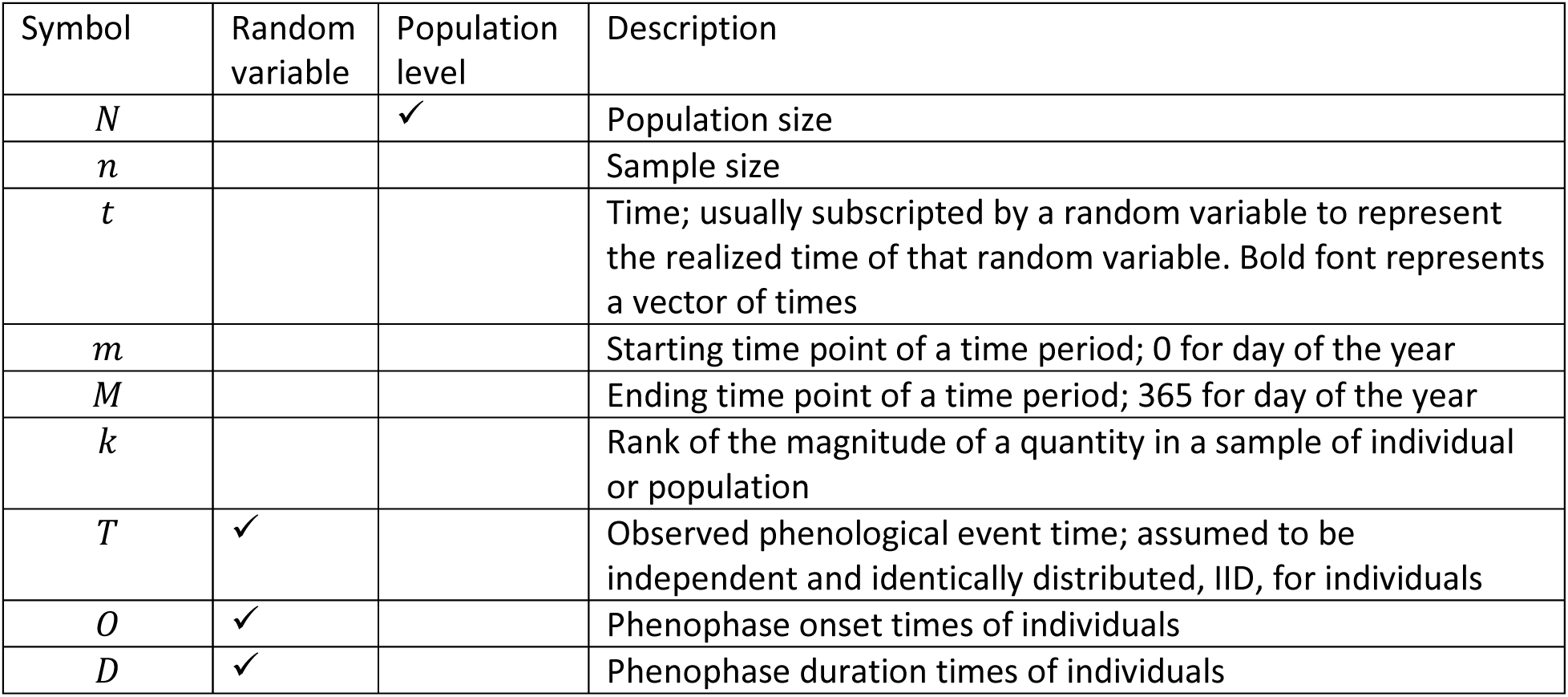
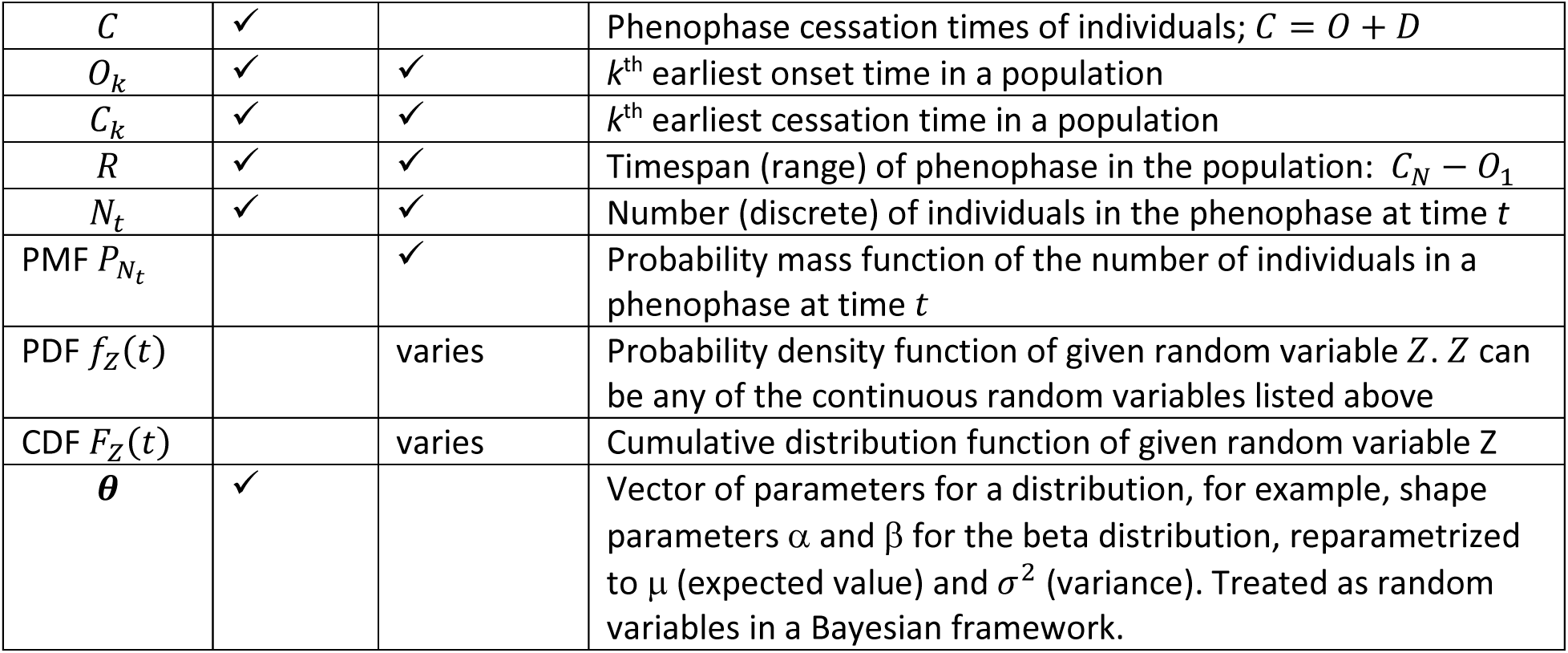
Listing of random variables, parameters, and their attributes.

| Symbol | Random variable | Population level | Description |
| --- | --- | --- | --- |
| $N$ | | ✓ | Population size |
| $n$ | | | Sample size |
| $t$ | | | Time; usually subscripted by a random variable to represent the realized time of that random variable. Bold font represents a vector of times |
| $m$ | | | Starting time point of a time period; 0 for day of the year |
| $M$ | | | Ending time point of a time period; 365 for day of the year |
| $k$ | | | Rank of the magnitude of a quantity in a sample of individual or population |
| $T$ | ✓ | | Observed phenological event time; assumed to be independent and identically distributed, IID, for individuals |
| $O$ | ✓ | | Phenophase onset times of individuals |
| $D$ | ✓ | | Phenophase duration times of individuals |
| $C$ | ✓ | | Phenophase cessation times of individuals; $C = O + D$ |
| $O_k$ | ✓ | ✓ | $k^{\text{th}}$ earliest onset time in a population |
| $C_k$ | ✓ | ✓ | $k^{\text{th}}$ earliest cessation time in a population |
| $R$ | ✓ | ✓ | Timespan (range) of phenophase in the population: $C_N - O_1$ |
| $N_t$ | ✓ | ✓ | Number (discrete) of individuals in the phenophase at time $t$ |
| PMF $P_{N_t}$ | | ✓ | Probability mass function of the number of individuals in a phenophase at time $t$ |
| PDF $f_Z(t)$ | | varies | Probability density function of given random variable $Z$. $Z$ can be any of the continuous random variables listed above |
| CDF $F_Z(t)$ | | varies | Cumulative distribution function of given random variable $Z$ |
| $\theta$ | ✓ | | Vector of parameters for a distribution, for example, shape parameters $\alpha$ and $\beta$ for the beta distribution, reparametrized to $\mu$ (expected value) and $\sigma^2$ (variance). Treated as random variables in a Bayesian framework. |

In probability theory, a RV is a mathematical mapping between a set of (often physical) states and a set of (real-valued) numbers that usually represent some type of measurement of the underlying states. Associated with those real-valued numbers is a probability distribution that defines the probability of obtaining a set of real-numbered outcomes. The onset RV *O* maps the (physical) time when an individual begins a phenophase to the (real-valued) time measurement that occurs within a repeating TP. The time measurement, such as day of year (DOY), is treated as a continuous variable, and when the year is the unit of the TP, DOY ranges between a minimum time 0 days (immediately after December 31^st^) and a maximum time 365 days (immediately before January 1^st^ on non-leap years). Associated with *O* is the probability that a phenophase will begin within a given set of days. This probability is modeled by the probability density function (PDF) or, more generally, by its cumulative distribution function (CDF). In the case of a continuous RV such as *O*, its PDF is represented by 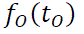. Its CDF is the probability that the onset is less than or equal to a given time: 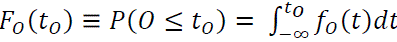.

### Derivations Overview

Phenological events, such as first onset, onset, peak phenophase, cessation, and last cessation, are hidden (latent) variables when biological specimens are used for inferences. None of these phenological events of interest are directly observed, and instead the observed collection time, hereafter referred to as the response variable, *T*, is used to infer the hidden variable states. A goal of this paper is a “reverse engineering” one: given randomly observed times when individuals are in a phenophase, infer the parameter values of the distribution of *T*. Then, given these parameter values, infer the parameters of the distributions of hidden variables.

Our mathematical derivations, however, follow a “forward engineering” approach.

Results of these derivations are summarized in Table 2. Our theory is built from models of the onset *O* and duration *D*. Ultimately, all other distributions, including the theoretical distribution of observed times *T*, are functions of the distributions for *O* and *D*. Consequently, when the theoretical distribution of observations is fit to empirical data, the onset and duration distribution parameter values are inferred, and the reverse engineering goal is achieved.

**Table 2.**
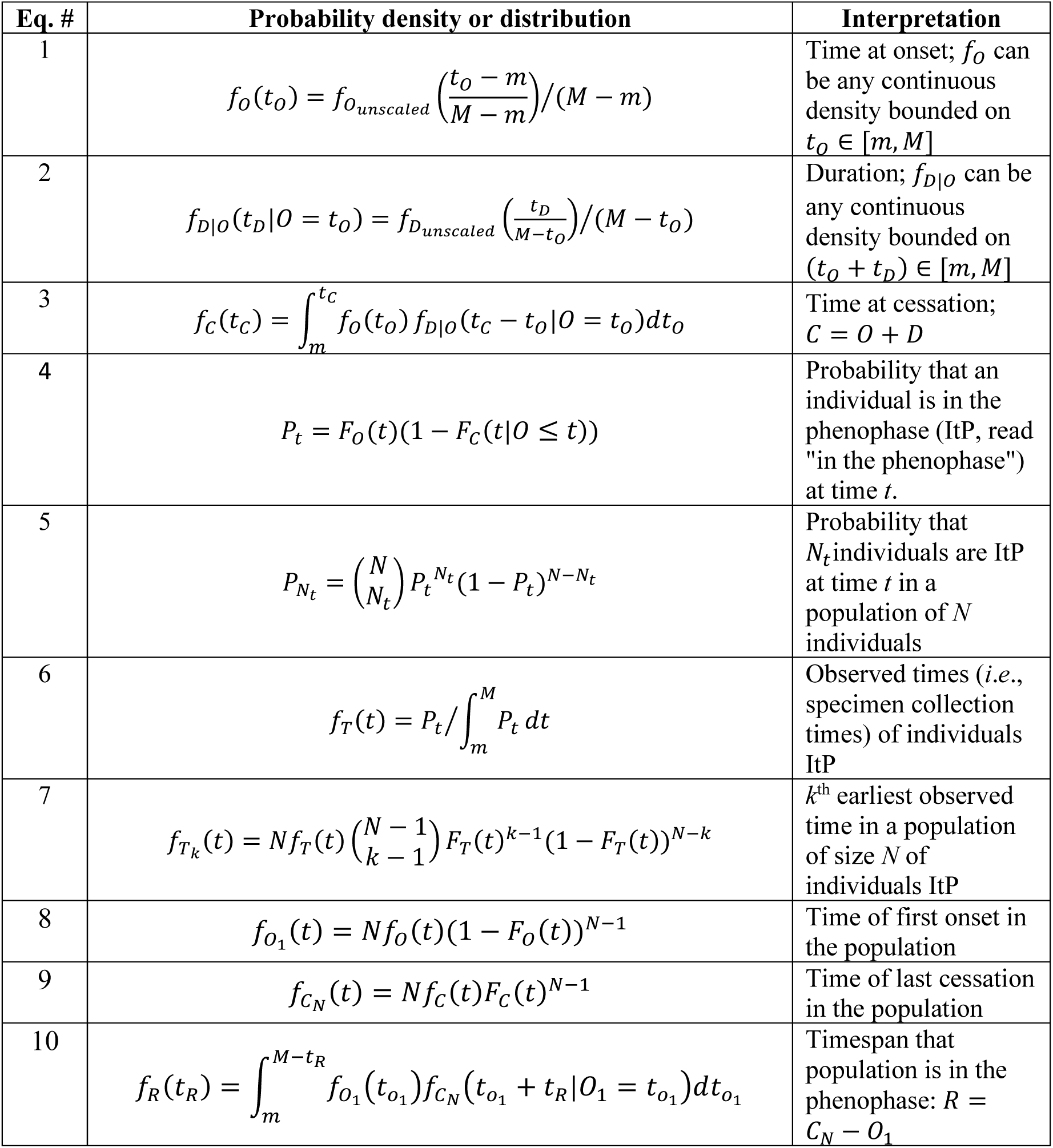
Unified theory of phenological timing.

### Onset (O), Duration (D), and Cessation (C)

The eight Properties place restrictions on which distributions are possible for *O* and *D*. Because phenophases are constrained to occur within finite TPs (Properties 2 and 3), the PDFs must have non-negative values within the TP and be zero outside the TP. Any distributions with upper and lower bounds on measured outcomes (i.e., finite support) could be used when appropriately scaled between the minimum (*m*) and maximum (*M*) times within the TP. Beta, truncated normal, and uniform distributions are examples.

If the PDF one selects to model the onset has support on [0,1], a change of variable for PDFs scales the PDF between *m* and *M* as follows:

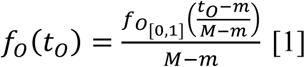

The duration, *D*, is dependent on the onset time, because the duration must be shorter than the interval of time between the onset time *t*_*O*_ and *M* (by Property 3). Starting from a PDF with support [0,1] for the duration, the scaled version given *O* = *t*_*O*_ is:

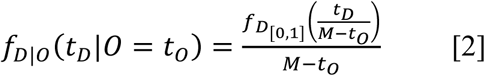

This, again, follows from a change of variable to scale the duration time *t*_*D*_ between zero and *M* − *t*_*O*_.

The sum of the onset time *O* and duration *D* determines the cessation time *C*: *C* = *O* + *D*. As a sum of random variables, the PDF associated with *C* is computed using the convolution formula, generalized to the case where the summands are not independent.:

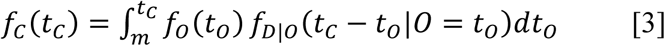

This result follows from the law of total probability (the “or” rule), the product rule of probability (the “and” rule), and the definition of conditional probabilities. Specifically, the probability of a cessation at time *t*_*C*_ can be found by summing (integrating for the continuous case) over all combinations of onset and duration times that satisfy *t*_*C*_ = *t*_*O*_ + *t*_*D*_. Let *A* represent the event that the onset time *O* falls in an interval *o* and *B* be the event that the duration time *D* falls in an interval *d* their joint probability is the probability that both events occur, *P*(*A*, *B*). By the definition of conditional probabilities, this equals *P*(*A*)*P*(*B*|*A*). *P*(*B*|*A*) is the conditional probability of *B* given *A*. In the continuous case, the probability mass function (PMF) *P* is replaced by the probability density function (PDF) *f* when single-point events like individual times (which have zero probability) are modeled.

### Peak Phenophase and the Distribution for Observed times (T)

Equations [1] – [3] define the population-level distributions for the onset, duration, and cessation times of a phenophase. The next step is to connect these distributions to the distribution of randomly observed times, modeled by *T*, of individuals that are in the phenophase of interest. First, equations [1] and [3] are used to determine the probability *P*_*t*_ that an individual selected at random from the population at time *t* is in the phenophase. Then, normalizing *P*_*t*_ defines the PDF for *T*.

For an observed time *t* to fall within the phenophase of an individual, it must occur after the onset of the phenophase (event *A’*) and before its cessation (event *B’*). The probability that *t* is after the onset is given by the complement of the CDF of onset times, 1 − *F*_*O*_(*t*) by the complement rule of probabilities. The probability that *t* is before or at the cessation time is given by the CDF of cessation times, *F*_*C*_(*t*). Combining these, the probability that *t* falls within the phenophase – i.e., after the onset *and* before the cessation – is:

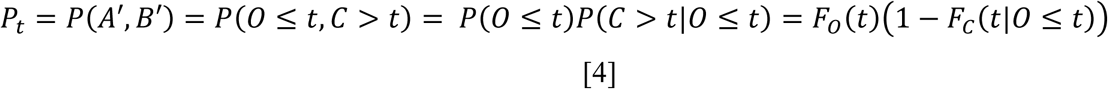

This expression captures the probability that an individual is in the phenophase at time *t*. The peak of the phenophase in the population corresponds to the time at which *P*_*t*_ is maximized, found by differentiating equation [4] with respect to *t*, setting the derivative equal to zero, and solving for *t*, or alternatively, by numerical optimization to find the value of *t* that maximizes *P*_*t*_.

Generalizing from peak phenophase, the probability distribution of the number of individuals *N*_*t*_ that are in the phenophase at time *t* is binomially distributed with parameters *P*_*t*_ and *N* (the size of the population):

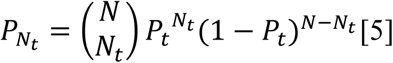

This distribution describes the likelihood of observing exactly *N*_*t*_ individuals in the phenophase at time *t*, given the total population size *N* and the time-specific phenophase probability *P*_*t*_. To express this in terms of percent-of-population, *N*_*t*_ is divided by *N* and multiplied by 100.

The PDF of the observed time *T* when specimens of individuals in the phenophase were collected can be obtained by normalizing the time-varying probabilities *P*_*t*_ over the interval of the time period [*m,M*]. This normalization ensures that the resulting function integrates to 1, satisfying the definition of a valid PDF:

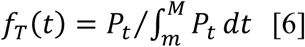

### Phenological Extremes

Phenological extremes refer to population-level events, such as first onset *O*_*k*=1_ ≡ *O*_1_, the earliest time at which any individual in a population initiates the phenophase, and the last cessation *C*_*N*_, the last time at which any individual concludes the phenophase. These extreme values are themselves random variables with associated PDFs, which model variation across multiple populations of individuals. For a population of *N* individuals, the PDF of the *k*^th^ earliest onset time is:

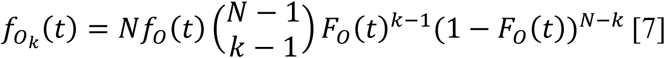

This expression follows from the standard formula (Mood, 1974, p. 253) for the PDF of the *k*th order statistic of IID random variables (Properties 6-8). Its derivation can be intuitively understood as follows. Imagine *N* slots each holding one individual from the population. The individual with the *k*th earliest time can occur in any of the *N* slots and occurs with “infinitesimal probability” *f*_*O*_(*t*). The remaining *k*-1 individuals with earlier times are binomially distributed. That is, there are 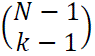 ways to choose where the remaining *k*-1 individuals with earlier times (each occurring with probability *F*_*O*_(*t*)) fit into the remaining *N-1* slots. Finally, the *N-k* individuals with later times (each occurring with probability 1 − *F*_*O*_(*t*)) fill the remaining slots. The product of these terms yields the PDF of the *k*th order statistic under the IID assumptions, equation [7].

The first onset time in the population is modeled by setting *k* to 1, resulting in the PDF:

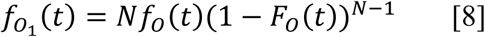

Replace the *O* in equation [8] with *C* to obtain the PDF of the *k*^th^ earliest cessation time in the population. Setting *k*=*N* provides the PDF of the last cessation time *C*_*N*_ in the population:

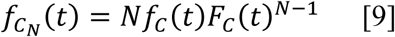

Equations [8] and [9] represent PDFs of phenological extremes in the population. Each population provides a single value of a phenological extreme. A *sample of populations* of the same size is needed to empirically estimate the mean earliest onset or last cessation. However, the theory (equations [8] and [9]), together with IID assumptions, allow the estimation of extremes with just the knowledge of parameter values that define the onset and cessation (or onset and duration) distributions of a single population.

The PDF of the total length of time during which at least one individual in a population is in a phenophase (i.e., the time range *t*_*R*_; *R* = *C*_*N*_ − *O*_1_) can be derived in parallel to equation [3] by considering all first onset times *t*_*O*1_ with last cessation times equal to *t*_*O*1_ + *t*_*R*_:

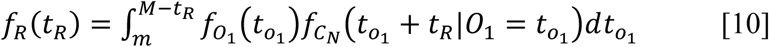

For normally distributed random variables *X* ∼δ(μ, σ), asymptotic approximations of the expected value E[*X*_*k*_] are available (Elfving, 1947; Cody, 1993)

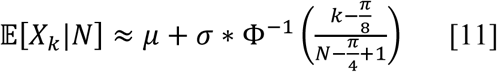

Φ^−1^(*x*) is the inverse of the CDF of the normal distribution.

### Statistical Properties of Phenological Distributions

Statistical properties of phenological distributions follow from standard definitions. Expected value, variance, and other moments of a RV can be calculated using the standard equations for these quantities. Frequentist 100(1-α)% probability intervals are calculated as 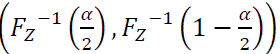, where 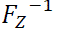 is the inverse of *Z*’s CDF, also known as the quantile function. The RV *Z* can be replaced by any of the RVs in Table 1.

### Distributional forms for Onset, Duration, and Cessation

The specific distribution used to model *O* or *D* is at the discretion of the researcher and can be tailored to the biology of the study system. Here, we consider two approaches. In the first, beta distributions are used for both *f*_*O*[0,1]_ and *f*_*D*[0,1]_ in equations [1] and [2]. The beta distribution is typically parameterized by two positive shape parameters, denoted α and β, that can be loosely interpreted as counts of successes and failures of a binomial process. The beta distribution can be reparametrized more intuitively by its expected value 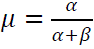 and its precision ϕ = α + β, or alternatively by its expected value and variance 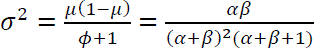. This parameterization enables intuitive control over the skewness, modality, and dispersion of the distribution, allowing for rich and flexible modeling of biological variation. Hereafter, we use the μ and σ parameterization, as α and β will be used to represent the intercepts and slope coefficients of linear models (to be described in the section *Modeling Covariates*). Thus, we can talk about the mean of the distribution of onset times as μ_*O*_ and its standard deviation as σ_*O*_. This subscripting convention will be used for all parameters and associated random variables. The beta-onset, beta-duration (BB) model therefore includes four parameters to be estimated by the data: the means μ and standard deviations σ for the onset and duration distributions. Two other parameters, the *m* and *M* bounds of the TP, are also needed, but these are known *a priori* and do not need to be estimated from the data.

The BB model obeys all eight Properties. However, after preliminary analyses, numerical computations involving differentiation and integration were prone to instability, and the computational cost to carry out a full Bayesian inference pipeline was prohibitively high with the BB model. To address these limitations, we explored a second, simplified modeling approach that, while violating some assumptions (notably Property 4), proved to be computationally feasible.

The second approach is also more biologically grounded in some respects. Rather than specifying ad hoc distributions like the beta distribution in the BB model, distributions for onset and cessation emerge from an explicit stochastic model of phenological signaling. Specifically, we model phenological timing as the result of a Gaussian process (GP; Rasmussen & Williams, 2008). Conceptually, this reflects a latent phenological signal – such as an endogenous physical cue or an exogenous environmental factor like temperature – that changes over time.

A Gaussian process defines a distribution over functions such that the functions’ values at any finite set of time points follow a multivariate normal distribution. We adopt a simple GP with linearly increasing mean and constant variance: 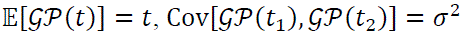 for all *t*_1_, *t*_2_ ∈ [*m*, *M*]. This implies that the process is homoscedastic with expected linear change in the signal over time. Under this model, we define the onset time as the first time the GP reaches a threshold μ_*O*_, resulting in *O*∼δ(μ_*O*_, σ). Similarly, the cessation time is modeled as the first time the GP reaches a second threshold μ_*C*_, yielding *C*∼δ(μ_*c*_, σ). Because both thresholds are crossed by the same realization of the process, the duration, *D* = *C* − *O*, is deterministic and equal to μ_*D*_ = μ_*C*_ − μ_*O*_. That is, the model does not allow for intrinsic developmental stochastic variation in duration. While this may seem restrictive, it reflects a biologically plausible scenario: under consistent environmental and physiological conditions, phenological durations are expected to be repeatable. Variation in duration still arises across individuals or populations due to differences in these conditions. In violation of Property 4, this GP model also places no bounds on when phenological events occur, which is not problematic when most of the density of phenological events occurs within a TP.

In summary, this GP-based model is a simplification of the BB model. It involves only a single standard deviation parameter σ, uses a normal distribution for onset and one with the same variance to model cessation and assumes no variability in duration (modeled as a Dirac delta distribution). Despite its simplifications, the GP model offers a biologically intuitive and computationally tractable alternative to more complex models.

### Modeling Covariates

The above framework provides a model of phenological events within and across populations but does not account for the influence of environmental or genetic factors. A primary goal of phenological analysis is to connect variation in phenological events to variation in underlying factors, which we refer to as covariates (to avoid implying causation). Let ***X*** = (*x*_1_, *x*_2_, …, *x*_*K*_) denote a vector of *K* covariates. To incorporate covariate effects on onset and duration, we model the expected onset time μ_*O*_ as a deterministic linear function of the covariates: 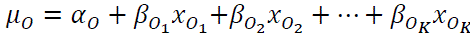, or more compactly in vector notation, 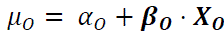. Similarly, the expected duration is modeled as 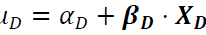 where covariates for duration ***X***_***d***_ may differ from those for onset. Under this setup, the full GP model has the following free parameters to be determined by data: one standard deviation parameter σ, two intercepts (α_*O*_, α_*D*_), *K*_*O*_ + *K*_*D*_ coefficient parameters (β_***o***_, β_***d***_). The two mean parameters (μ_*O*_, μ_*D*_) are fully determined by the intercepts and coefficients.

Consequently, under the GP process,

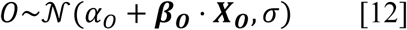

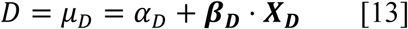

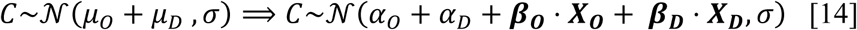

When the same covariates ***X*** are used to model onset and duration, this simplifies to: 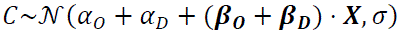. Thus, 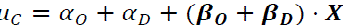. In this paper, we will use the same covariates to model onset and duration.

Since *O* and *C* are both normally distributed in the GP model, phenological extremes can be approximated by equation [11]. Plugging in the linear models for the means gives the models for the expected extremes in terms of the covariates: 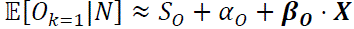 and 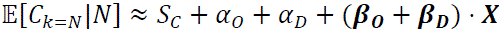, where 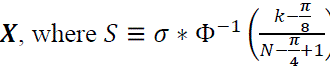 is constant for a population of size *N*. These expressions give computationally-efficient approximations for phenological extremes.

When biological specimens are collected randomly during the phenophase (Property 8), *T* has a symmetrical distribution under the GP. The expected observed time is midway between onset and cessation for symmetrical distributions: 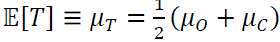. Substituting expressions for μ_*O*_ and μ_c_ gives 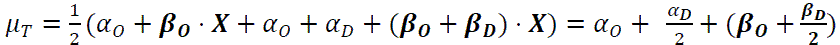 Since the least squares regression line goes through the mean values, this expression also describes the regression model for observed collection times as a function of covariates under a standard linear regression model (SR).

However, performing SR will not allow the individual contributions of onset and duration covariates to be disentangled due to perfect collinearity between β_***o***_ and β_***d***_ in their combined effect on *T*. Despite this limitation, SR provides a useful reference point for evaluating the performance of a full Bayesian analysis. In particular, the intercept 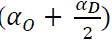 and slope coefficients 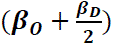 derived from the parameters of the GP model can be directly compared to those obtained from SR when fit to the observed times. When a Bayesian analysis has performed well, these quantities should closely match the SR estimates, serving as a consistency check on the model’s inference. They are not expected to be equivalent because the prior distribution can bias inferences.

### Bayesian Inference

Now that the theory to model the observed times is available (given by equations [6], [12-14]), the next task is to select a procedure to infer the parameter values for *T* from empirical data. Two approaches to probabilistic inference are common. Maximum likelihood estimation (MLE) finds parameter values that maximize the probability of the observed data given the model and parameter values. The likelihood function is written as a function of the vector of parameters θ whose values are to be estimated. The following likelihood function results from the IID assumptions (Properties 6-8) applied to equation [6]. Here, ***t*** = (*t*_1_, *t*_2_, …, *t*_*n*_) represents a vector of *n* observed collection times:

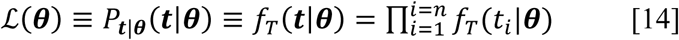

MLE provides point estimates of parameter values. When a goal is also to estimate the uncertainty associated with parameter estimates, a Bayesian approach can be used. Bayesian inference treats parameters as RVs with associated PDFs and makes it possible to ask what the probability of the true parameter value of the population is given the data and the model. This Bayesian density, called the posterior *P*_θ|***t***_(θ|***t***), is the conditional probability density of the parameter values given the observed times ***tt*** and given the model specification (e.g., GP). The posterior is provided by Bayes rule:

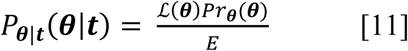

The posterior distribution is a function of the likelihood and the prior 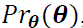, scaled by the reciprocal of the constant evidence value *E*, which assures normalization of the posterior. The prior specifies the PDF of parameters based on a prior understanding. When new observations become available, the posterior distribution from a previous analysis can successively be used as the prior in subsequent analyses. In such a way, a Bayesian analysis can iteratively improve one’s understanding of phenomena as new data become available and is highlighted as the unique statistical approach that does so validly by Jaynes (Jaynes, 2021). We apply this iterative Bayesian approach to phenological estimation as described in section *Analysis of the Phenology Paradox using Bayesian Inference, Quantile Regression, and Standard Linear Regression*.

Using Monte Carlo techniques, it is not necessary to calculate the normalizing constant *E* ≡ *P*(***t***), as it cancels out during the inference process. Nevertheless, understanding how it is calculated is instructive because the procedure – marginalization – is central to the Bayesian framework that underlies our approach to simulations and analyses. Marginalization enables the computation of the distribution of a variable of interest *T* by integrating over the uncertainty or variability associated with another variable, θ, that provides information about *T*. Marginalization is expressed as 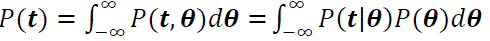 where *P*(***t***|θ) is the likelihood function (i.e., equation [14]), and *P*(θ) is the prior distribution over θ (i.e., 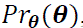). In the context of Bayesian inference, marginalization plays a central role by allowing one to integrate over uncertain or latent parameters to obtain the marginal distribution of an observable variable (e.g., *P*(***t***)) or to compute expectations when summarized across uncertain conditions. Marginalization thus provides a principled way to isolate the distribution of a single component of a composite quantity by appropriately accounting for the uncertainty in its underlying components.

In general, a closed-form of the posterior distribution is unavailable, and its properties are approximated by Monte Carlo techniques, such as Metropolis-Hastings Markov Chain Monte Carlo (MH MCMC) (Metropolis *et al*., 1953; Hastings, 1970), which produce a valid sample of parameter values from a posterior distribution. We implemented our GP model in the probabilistic programming language, Stan (Carpenter *et al*., 2017), which uses Hamiltonian Monte Carlo (HMC) (Neal, 2011; Betancourt & Girolami, 2013) to carry out the sampling from the posterior distribution. We refer to this inference procedure as Bayesian GP.

### Proof of the Phenology Paradox

Here, we demonstrate mathematically that under certain combinations of parameter values, simple linear regression (SR) models, when fitted to observed collection times, are guaranteed to fail in the large-sample limit when their slope coefficients are interpreted as proxies for onset phenological sensitivity. The coefficients of SR models fit to observed times are composites of coefficients from both onset and duration models. Specifically, the relationship between the regression coefficients β_***T***_ of the SR model, and the coefficients for onset and duration, is given by 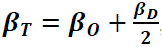, as derived in *Modeling Covariates*. This identity illustrates that β_***o***_ (onset) and β_***d***_ (duration) contribute to the observed-time slope β_***T***_. SR models cannot disentangle these two components. When one component dominates the other, the weaker signal may be obscured entirely in β_***T***_. There are six feasible sign combinations for β_*O*_, β_*D*_, and β_*T*_ for a single covariate, illustrated in Fig. 2. Two of these scenarios are particularly problematic. We call them paradoxical because the sign of the onset model slope β_*O*_ is opposite to that of the observed-time slope β_*T*_. In such cases, using β_*T*_ as a surrogate for β_*O*_ not only produces inaccurate estimates, but the direction of the inferred effect is wrong.

**Fig. 2.**
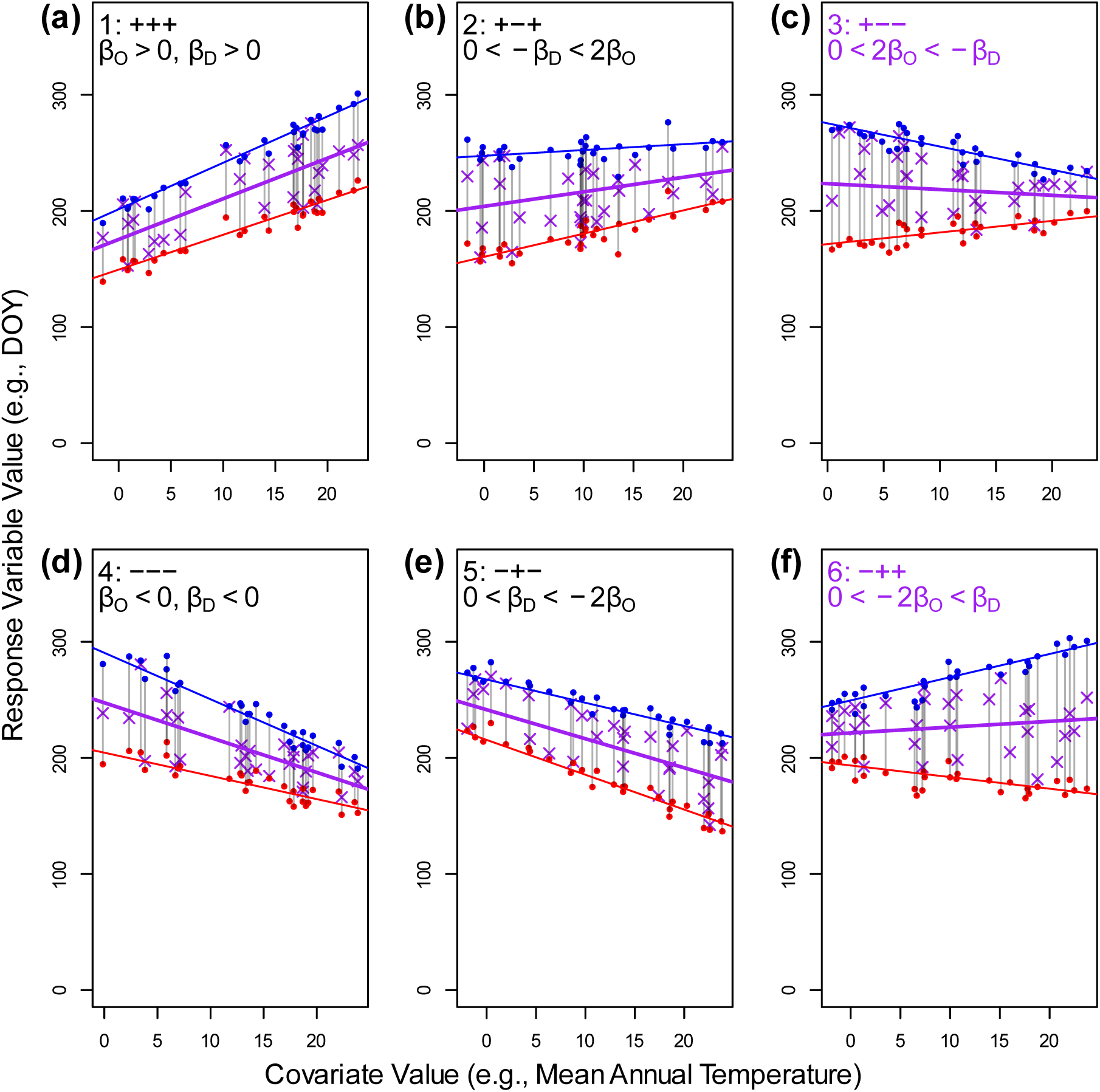
Conditions giving rise to six Scenarios, including two paradoxical Scenarios. Each scenario is defined by a triplet of plus or minus signs indicating the direction (positive or negative) of the regression slopes for onset (β_O_) and duration (β_D_) and observed-time (β_T_) models, respectively. Inequalities indicate conditions when the third plus or minus sign in each triplet (that represents β_T_’s sign) is achieved. As in Fig. 1, each line segment represents an individual experiencing a phenophase, beginning at the red dot (onset) and ending at the blue dot (cessation); the segment length corresponds to the duration. Purple x’s indicate the time of observation. The *x-*axis of Fig. 1 is rotated here and serves as the *y-*axis. Individuals are sampled at different values of a covariate shown on the *x*-axis (e.g., temperature or precipitation). In paradoxical Scenario 3, a large decrease in duration offsets increasing onset values; in Scenario 6, a large increase in duration offsets decreasing onset. Data for these graphs came from an arbitrarily selected set of simulation replicates from the “Phenology Paradox” simulations.

For example, in Scenario 3, we might observe a negative β_*T*_ for a latitude covariate suggesting that phenological events occur earlier at higher latitudes. However, the true onset slope β_*O*_ would be positive in Scenario 3, indicating that onsets actually occur later with increasing latitude. This scenario arises whenever the negative term 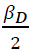 outweighs the positive signal from β_*O*_, resulting in a negative β_*T*_. Paradoxical Scenario 6 occurs under the reverse conditions: 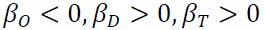. In this case, the positive contribution from 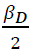 overwhelms the negative signal from β_*O*_, resulting in a misleading positive β_*T*_. A full enumeration of all six Scenarios and their mathematical conditions is provided in Fig. 2. We call these outcomes the Phenology Paradox.

In contrast, when duration is independent from the covariate, β_*D*_ is zero, so β_*T*_ = β_*O*_. In this case when duration doesn’t change systematically with changes in the covariate, the slope of the SR model fit to the observed times will equal the slope of the onset model.

### Model Validation

As a completely new model, we carried out multiple analyses to assess the strengths and limitations of our approach, including an (1) assessment of the match between theoretical distributions (Table 2) and simulated observations, (2) accuracy of Bayesian GP when the observations were generated under a different model than the GP (i.e., model mis-specification), (3) issues of model identifiability and prior distributions, and (4) comparisons to the results of previous methods. In particular, we compare our approach to Pearse et al.’s estimator (PE; (Pearse *et al*., 2017)) to analyze phenological extremes, and we compare our results to both Park et al.’s quantile regression approach (QR; (Park *et al*., 2025)) and to SR for onset, peak observed, and cessation events. We end our analyses with Bayesian GP applied to empirical data sets.

### Match Between Theory and Simulation

Ideally, empirical data about the true onset, duration, and cessation times based on careful field observations of individuals in populations would be available to evaluate the effectiveness of the proposed Bayesian GP method to compare results to known empirical quantities. Although such data sets are available for some species (e.g., Davis *et al*., 2015; Renner *et al*., 2021), such data sets are relatively scarce, in general, although they are increasing in frequency due to the efforts of organizations such as the National Phenology Network (Crimmins, 2021) and teams of researchers who have gathered extensive datasets of field observations for hundreds of species that span multiple centuries (Amano *et al*., 2010; Büntgen *et al*., 2022).

We used simulated data based on known parameter values to evaluate the error between inferred parameters and the true parameters used to simulate the data. For this approach to work, we first verified that the properties of the simulated data closely match the properties of the theoretical distributions. We tested this correspondence using the Kolmogorov-Smirnov test as implemented by the R *stats* package function `ks.test` for 1,000 sets of randomly chosen, but feasible parameter values (i.e. resulting in distributions falling within a year TP) for the onset and duration distributions for both the BB and GP models. For all simulations, *m* and *M* were set to 0 and 365, respectively, corresponding to the minimum and maximum day of year (DOY) for a year time period. Full simulation parameter settings are recorded in Table S1 under the simulation “Monte Carlo Simulation / Theory”.

Simulations for the onset were straightforward. For BB, the simulation mean and standard deviation parameters were scaled between 0 and 1 according to the scaling provided by equations [1] and [2]. For BB, onset times were drawn from the beta distribution using R’s `rbeta` function or from the normal distribution using `rnorm` for GP. Cessation times under GP were found by adding the constant μ_*D*_to the sampled onset times. Duration for BB was again sampled using `rbeta`, then values were scaled according to equation [2]. Simulated values for BB were rescaled back to the original scale by multiplying by *M*-*m* (365-0). To simulate observed times, *T*, a random value between a simulated onset time and cessation time was uniformly sampled using `runif.` First onset and last cessation times were simulated by first simulating the onset and cessation times for a population of *N* simulated individuals. The first onset time was the earliest simulated onset time in the population, and the last cessation time was the largest cessation time in the population. To obtain a Monte Carlo distribution of these phenological extremes, 1,000 populations each of *N*=1,000 were simulated. Finally, the population phenology range (equation [10]) was simulated as the difference between the first onset and last cessation for each of the 1,000 simulated populations.

### Model Mis-specification

Additional simulations were then used to evaluate how well inference using Bayesian GP recovered the true parameter values. Table S1 “Validation” provides full details of ranges of parameter values, hyperparameters, and sample sizes used during the simulations. Values were uniformly randomly sampled between the ‘Minimum’ and ‘Maximum’ row values in the correspondingly-labeled columns of Table S1. Raw error and absolute error were recorded for each model parameter for each simulation replicate. The mean error estimates were marginal quantities since they were averaged across variation in sample size, hyperparameters, and model parameters. Differences in error rates were tested using the *t*-test implemented by R’s `t.test` function.

### Model Identifiability and Priors

A model is identifiable when a unique combination of parameter values defines the shape of the distribution. When multiple combinations of parameter values result in the same shape, the model is non-identifiable. In this study, inferences were made from the distribution of observed collection times *T*. If distinct combinations of onset and duration parameters result in similarly-shaped observed-time distributions, then non-identifiability can arise, potentially leading to inferences that fail to converge to the true parameter values. In Bayesian inference, priors act as regularizers that penalize implausible regions of parameter space (Bishop, 2006). By incorporating prior knowledge or constraints, they can thereby improve convergence and stability of posterior sampling, particularly in the presence of low-information or ambiguous data. In contrast, maximum likelihood estimation (MLE) does not use priors, relying solely on the likelihood. As such, if identifiability issues are present, they are more likely to manifest clearly during MLE, making it a useful diagnostic tool for detecting model identifiability issues. MLE was therefore also used to infer parameter values from the simulated data.

We also used the simulated “Validation” data set to explore the impact of biases in hyperparameter values used during inference. Hyperparameter bias was defined as the difference between the hyperparameter value used during the Bayesian analysis and the true parameter value. We hypothesized that inference error would be positively correlated with hyperparameter bias, but that the presence of informative data would mitigate this error by guiding the inference process closer to the true values. We examined this hypothesis using linear regression, implemented by the R `lm` function.

### Comparison to Pearse Estimator (PE) of Phenological Extremes

To explore the efficacy of our approach to estimate phenological extremes, we compare Bayesian GP estimates to those based on the Pearse et al. (Pearse *et al*., 2017) estimator (PE) Pearse et al. recognized that the distribution of phenological extremes is well approximated by the Weibull distribution. They therefore applied an approach based on this theoretical result to estimate phenological extremes, specifically, first onset events and claimed that their method works for any underlying distribution of data.

Based on our theory, once the onset and duration parameters are inferred from the data, these parameters can be used to estimate the timing of extreme phenological events modeled by equations [8], [9], and [11]. Using an additional set of simulations under GP, we compared raw mean error between estimates obtained by PE and estimates obtained by Bayesian GP by varying sample sizes, population sizes, and hyperparameter values. In contrast to the previous simulations, population sizes were set to fixed values: 100, 10,000, and 1,000,000 and sample sizes were deterministically set to 10 up to 90 by increments of 2. For each combination of population size and sample size, 1,000 simulation replicates were made. Full simulation specifications are provided in Table S1 “Pearse Comparison”. Since Pearse et al. (Pearse *et al*., 2017) claim their method is robust to any underlying distribution, simulation under the GP model should not bias inferences against PE.

Unlike PE, our method requires an estimate of population size, which was taken as given for both PE and Bayesian GP. Although, for PE, the population size has no effect on inference.

### Analysis of the Phenology Paradox using Bayesian Inference, Quantile Regression, and Standard Linear Regression

To examine the predictions of the Phenology Paradox, we carried out another set of simulations that incorporated a single covariate. These simulations also explore the accuracy of Bayesian GP when covariates are included in the model. Previous simulations above did not include covariates. The value of the covariate was randomly sampled between a minimum value (set to –2) and a maximum value (set to 24) to model variation in, e.g., mean temperature values across a large latitudinal gradient. Slope and intercept values were randomly sampled under the constraints imposed by the six Scenarios.

Once the slope parameter values for the onset and duration models were randomly obtained (subject to the constraints of the Scenario being simulated), and the covariate value was randomly sampled, the linear models for onset and for duration were applied to determine the mean for onset and duration. Sigma was held constant for these simulations since the goal of the analyses was to investigate the impact of varying slopes, not varying sigmas. With means and sigma in hand, simulations of observed times proceeded as described in *Match Between Theory and Simulation*. For each of the six Scenarios, we simulated 1,000 replicates.

We reparameterized the regression model in terms of an anchor rather than an intercept. The anchor can be thought of as the mean response (i.e., DOY) when marginalized across values of the covariate. In particular, the anchor, *A*, is related to the slope and intercept of the standard formulation as follows: 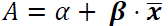 where 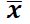 is the vector of covariate means. Since covariate values were randomly sampled between the covariate minimum and maximum values, 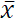 was one half the difference between the maximum and minimum. Full simulation details are provided in Table S1 “Phenology Paradox”.

This reparameterization in terms of the anchor was also used for Bayesian GP, as it improved posterior sampling diagnostics and made setting the prior more intuitive. It was not intuitive to set a prior on an intercept parameter, whereas the mean onset time, for example, is more intuitive.

Estimates of slope parameter values and anchor values were obtained using three approaches: Bayesian GP, quantile regression (QR), and standard linear regression (SR). In the SR approach, the slope of the regression line fit to simulated observed times was used as a proxy for the onset slope as has been done previously in the literature. Quantile regression was implemented following the recent methodology of (Park *et al*., 2025), who “arbitrarily” (their wording) assigned phenological phases to specific quantiles of the observed-time distribution *T*: the 10% quantile for onset, the 50% quantile for peak phenophase, and the 90% quantile for cessation. Accordingly, regression parameters estimated at the 10%, 50%, and 90% quantiles were interpreted as corresponding to onset, observed-time, and cessation distributions, respectively. Because the simulations were generated under a Gaussian Process (GP) model, the distribution of observed times is symmetric, such that the median (50% quantile) coincides with the mean, through which the SR regression line passes.

To set hyperparameter values for Bayesian GP, the simulated data were partitioned into two subsets: one used for hyperparameter estimation (30% of the data), and the other reserved for the Bayesian GP analysis (70% of the data). Quantile regression (QR) was applied to the first subset, and the resulting estimates were used to set the mean slope and anchor hyperparameters for the onset and cessation models for the Bayesian GP analysis. This approach is justified by a core principle of Bayesian analysis: using prior results (in this case, QR estimates) to inform the prior distribution hyperparameters. Because the data subset used to estimate hyperparameters was independent of the subset used by Bayesian GP, this strategy does not violate Bayesian principles about reusing data (which results in biased inferences). This approach parallels the training/testing framework commonly applied in supervised learning in the machine learning literature, where a model is trained on one subset and evaluated on another to prevent overfitting and assess generalizability.

Error was assessed in two ways. First the mean absolute error, i.e., the absolute value of the difference between the estimated and the true parameter value, was assessed. Second, the proportion of simulation replicates that correctly identified the sign of the onset, duration, and observed-time model slopes was recorded.

### Analyses of Empirical Datasets

Our final set of analyses used biocollection data of thirteen spring ephemeral flowering species native to northeastern North America. The primary objective was to evaluate how frequently phenophase duration is correlated with covariates. As shown above, when duration is independent of covariates, the slope of the SR model when fit to observed times provides a valid estimate of the onset slope. However, when duration varies systematically with covariates, SR slopes no longer reliably reflect onset dynamics. To our knowledge, explicit estimation of duration–covariate relationships using biocollection data has not previously been conducted. As such, this analysis provides important insight into the need for revised statistical approaches— such as those developed here—that account for duration effects when estimating phenological sensitivities.

### Data

Georeferenced specimen data and images were retrieved from iDigBio (https://www.idigbio.org/) via the *ridigbio* package (Michonneau *et al*., 2016) for each of the 13 species. Images associated with each specimen were manually screened to include only flowering specimens in subsequent analyses. Five covariates were used in the analyses: Year (as separate from DOY), mean annual temperature (obtained by averaging across mean monthly temperatures), mean spring temperature (obtained by averaging across monthly averages for March, April, and May), Elevation, and Latitude. Climate data were obtained from Berkeley Earth High-Resolution (Beta), which is an evolution of previous work (Rohde & Hausfather, 2020). This data set has monthly mean and maximum monthly terrestrial temperature values from the mid 1700’s up to “recent” (Berkely Earth language) at a spatial resolution of 0.25° X 0.25°. Temperature data were parsed using tools in the *ncdf4* R package (Pierce, 2025). Elevation values were obtained using the R package *elevatr* (Hollister *et al*., 2023). Response variable (DOY) and covariate outlier data were removed using Mahalanobis distance (Aggarwal, 2017; Li *et al*., 2019), studentized residuals (Williams, 1987), and Cook’s Distance (Cook, 1977; Cook & Weisberg, 1995). All incomplete records and records with duplicated data were removed.

Prior distribution hyperparameters were set to QR estimates using a partition consisting of 30% of the specimens, and 70% of the specimens were used for Bayesian GP. HMC sampling was done at default levels in Stan: 1,000 warm-up samples and 1,000 retained samples for each of 4 chains. R-hat values, divergence percentage, effective sample size (ESS), and E-BFMI diagnostics were examined to assess the quality of the sampling based on recommendations by Stan programmers, e.g. (Betancourt, 2017). R-hat compares between-chain and within-chain variances; successful runs have an R-hat close to 1. E-BFMI is a measurement of sampling efficiency and should be above 0.3. Divergences represent Markov Chain transitions that don’t follow the geometry of the posterior distribution. Divergences should be zero. ESS gives an indication of the effective number of independent draws from the posterior distribution. ESS should be in the high hundreds or above for the default sampling settings.

Results of the analyses of empirical data were summarized in tabular and graphical form. Marginal posterior probabilities for the sign of slope parameters were estimated as the proportion of samples of a slope parameter from the posterior distribution with a negative sign.

We also produced posterior predictive plots to visualize predicted observations 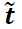 given previous empirical observations ***t***. The posterior predictive distribution quantifies uncertainty in future observations by marginalizing over the uncertainty in both the model parameters and covariate values (which employs the chain rule for probabilities):

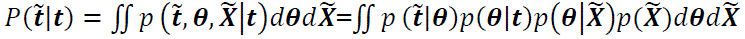

This integral is typically evaluated using Monte Carlo sampling. In our case, covariate values 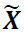 were first sampled from a joint distribution 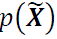, which captures empirical covariation among covariates. To approximate 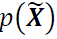, we used a Gaussian copula (Rank, 2007), which enables flexible multivariate modeling by coupling marginal distributions through a correlation structure.

We fit the copula model using the R package *copula* version 1.1-6 (Hofert *et al*., 2025). For each Monte Carlo draw, (1) a sample of 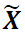 was drawn from the copula, (2) a draw of model parameters θ (slope, anchor, and sigma parameters) was taken from the posterior distribution obtained from the Stan run, (3) using the posterior draw of slope and anchor parameters together with the sampled covariate data, the μ parameters were determined using the linear models, (4) using the calculated μ parameters and sigma, new values of 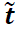 were simulated using the generative model 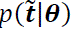 as described in the section *Match Between Theory and Simulation*. Posterior predictive summaries (e.g., the means and 95%CIs) were then computed for simulated 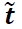. Finally, to smooth over random variation from Monte Carlo draws and copula artifacts, we smoothed the mean of simulated values using a generalized additive model (GAM) with the R package *mgcv*’s `gam` function with default options. The result was visualized to display the outcome of the Bayesian GP inference. For each value of a target covariate (represented by the graph’s x-axis), 1,000 Monte Carlo draws were made. This process allowed us to quantify uncertainty in the predicted values across the covariate range, reflecting both parameter uncertainty and covariate-driven variation.

## Results

Because this is a methods paper, we discuss important technical methodological considerations here and reserve the **Discussion** section to summarize the main outcomes and broader ramifications of this work.

### Match Between Theory and Simulation

The theory we derived aligns closely with data generated via Monte Carlo simulations. Using the Kolmogorov–Smirnov (KS) test, we evaluated whether the simulated distributions deviated significantly from theoretical expectations across 1,000 replicates. The proportion of replicates that rejected the null hypothesis at the α=0.05\alpha = 0.05α=0.05 level did not differ significantly from 0.05 in any case, as assessed using a binomial sign test (*O*: 39, p-value=0.13; *D*: NA – duration has no variance under the GP model; *C*: 48, p-value=0.83; *O*_*k*=1_: 58, p-value=0.25; *C*_*k*=*N*_: 58, p-value=25; *T*: 45, p-value=0.51). Figure 3 illustrates an example of a “pathological” case (defined later) showing the close agreement between simulated and theoretical distributions. These results support the use of Monte Carlo simulations as a practical and accurate alternative to more computationally intensive theoretical approaches.

**Fig. 3.**
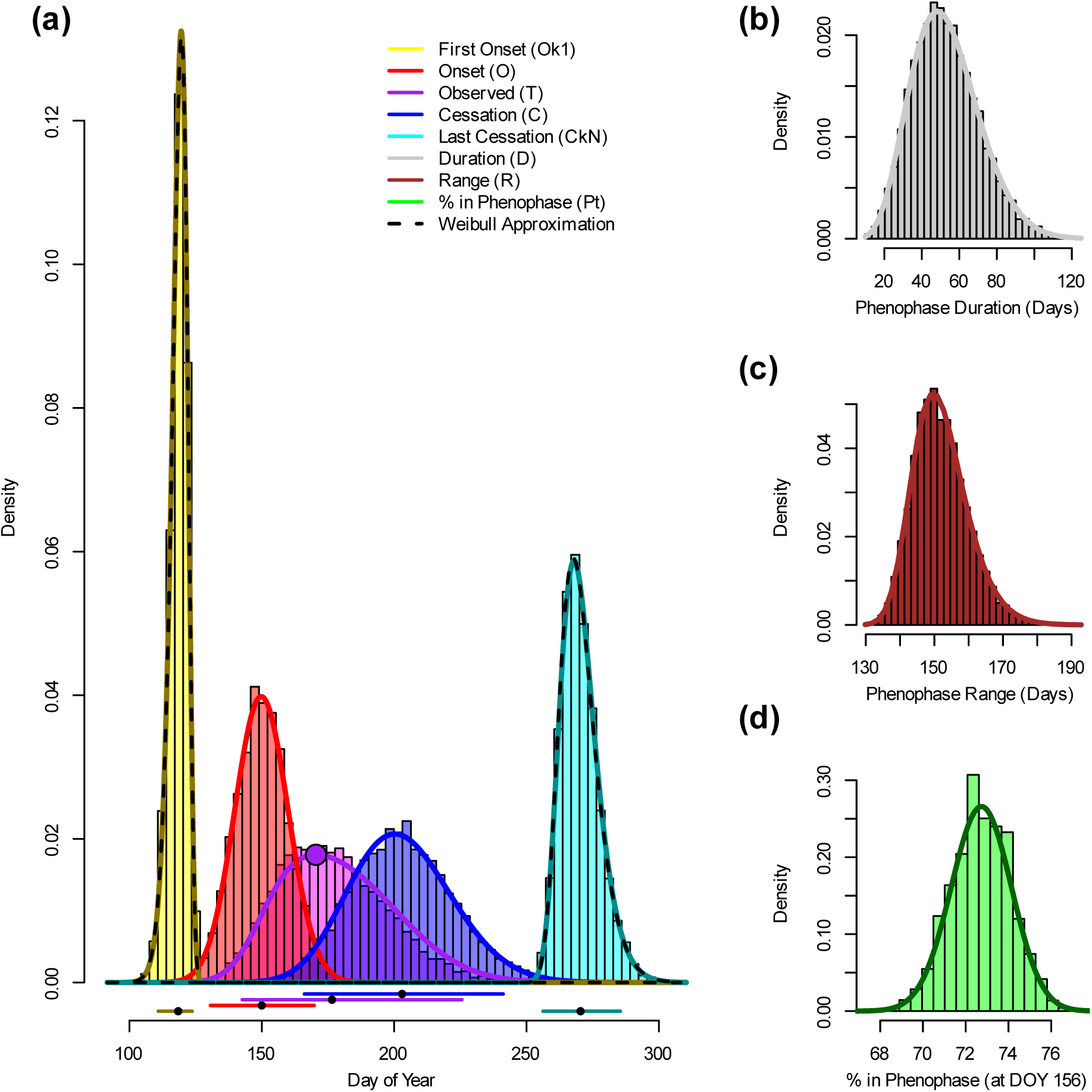
Agreement between theory and simulation across multiple phenological distributions. Simulation parameters were chosen to produce a skewed distribution of observed collection times *T*: μₒ = 150, µ_D_ = 90, σₒ = 10, σ_D_ = 30, and N = 1,000. Histograms are based on 10,000 simulated draws from these distributions, while solid curves represent the corresponding theoretical probability density functions (PDFs). (a) Core phenological distributions: first onset, onset, observed, cessation, and last cessation. The large purple dot indicates the peak phenophase at the population level. Horizontal intervals below the PDFs represent the mean (black dot) ± two standard deviations of these distributions. Dotted lines show Weibull distributions fit to the first onset and last cessation times. (b) PDF of individual phenophase durations. (c) PDF of total phenophase duration across all individuals in the population. (d) PDF for the probability that a given proportion of individuals (*x*-axis) is in the phenophase on a specific day of year (DOY), here arbitrarily set to day 156.

### Model Mis-specification

For all parameters, using the correctly specified model yielded lower mean absolute error (top number shown in each panel). However, the difference in mean absolute error between the BB and GP models was less than one day in all cases except for phenological extremes.

Differences in mean raw error (bottom number in each panel of Fig. 4) were also less than one day in all comparisons. So, inference under the mis-specified GP model had little practical impact on error metrics (Fig. 4). Error levels in Fig. 4 show values marginalized over simulated variation in sample size, model parameters, and hyperparameters, indicating expected outcomes in a general-use case scenario. For core μ parameters (μ_*O*_, μ_*C*_, μ_*D*_, μ_*T*_; Fig. 4a, d, f)) there was no statistical difference (*t*-test p-values all under 0.05) between mean absolute error when comparing accuracies of inferences from data simulated under the BB model vs. under the GP model.

**Fig. 4.**
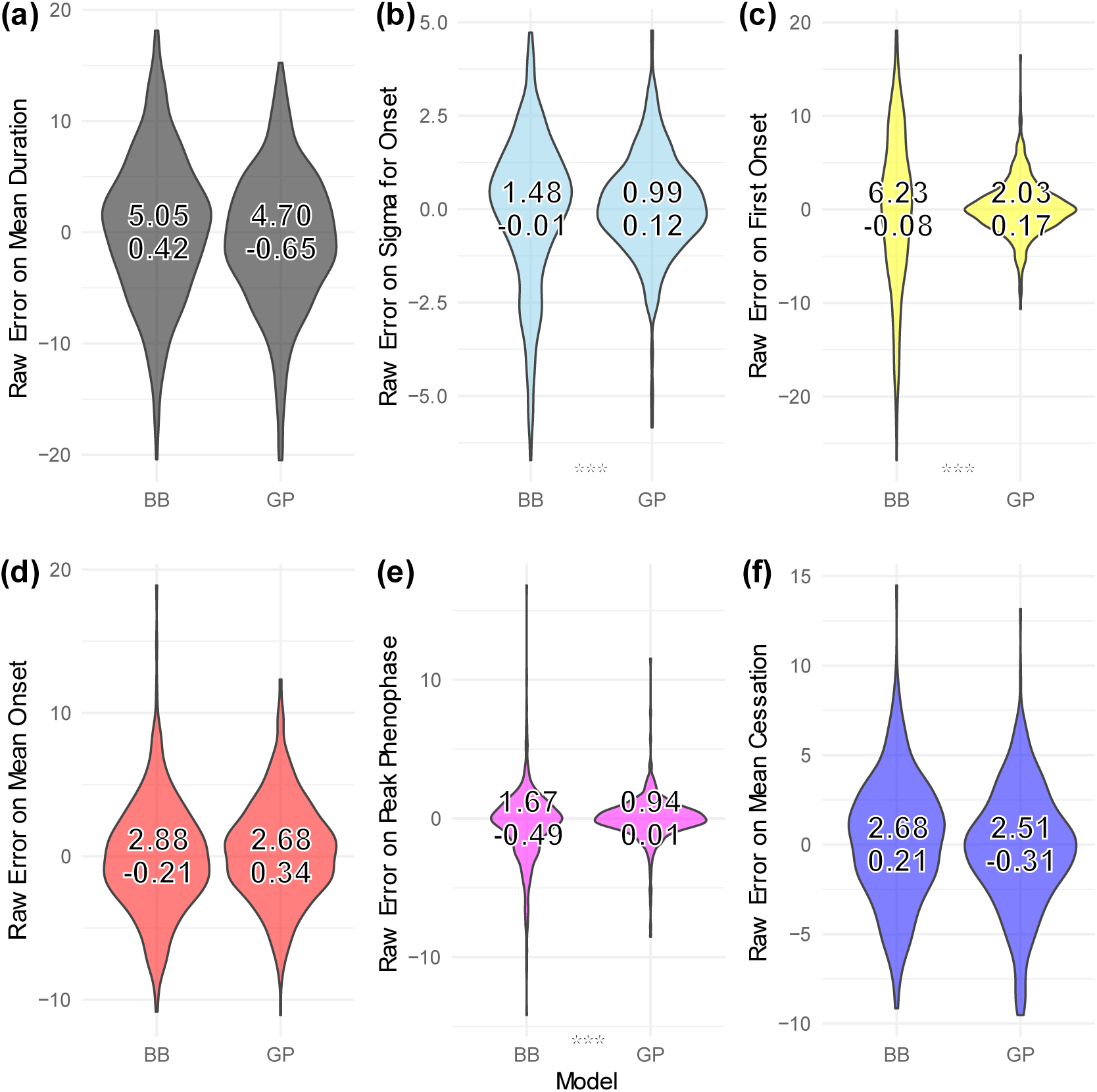
Model mis-specification. Panels (a)–(f) show violin plots of raw errors for the parameter estimates indicated on the y-axis labels. Data were simulated under either the beta-onset, beta-duration (BB) model or the Gaussian process (GP) model. In all cases, Bayesian GP inferences were made, regardless of the true data-generating process. Errors are marginalized over variation in prior distribution hyperparameters, sample sizes, and simulation parameter values, based on 1,000 simulation replicates. Mean absolute error (top number) ranged from less than one day (for peak phenophase under the GP model, (e)) to over six days (for first onset under the BB model (c)); bottom number is mean raw error. Asterisks (‘***’) denote statistically significant differences in absolute error at the α = 0.001 level.

In contrast, there were statistically different error levels between BB and GP for σσ, phenological extremes, and peak phenophase (Fig. 3b, c, e), but these differences were of little practical significance (less than a day, except for phenological extremes.)

Figure 5 illustrates a particularly pathological case in which the Bayesian GP performs worse. However, even in this extreme case, the true values of core μ parameters fall in the 95% CI. Because the true onset and cessation distributions have markedly different variances, the resulting observed-time *T* distribution is skewed. The GP model, however, assumes a shared variance parameter for both onset and cessation, and thus cannot accurately recover their individual variances. This mis-specification in variance propagates to larger errors in the inferred distributions for phenological extremes, as the expected extreme values depend on the product of the standard deviation and a function of population size (equation [11]). These considerations explain why there were statistically different error rates for the σ parameter and extremes.

**Fig. 5.**
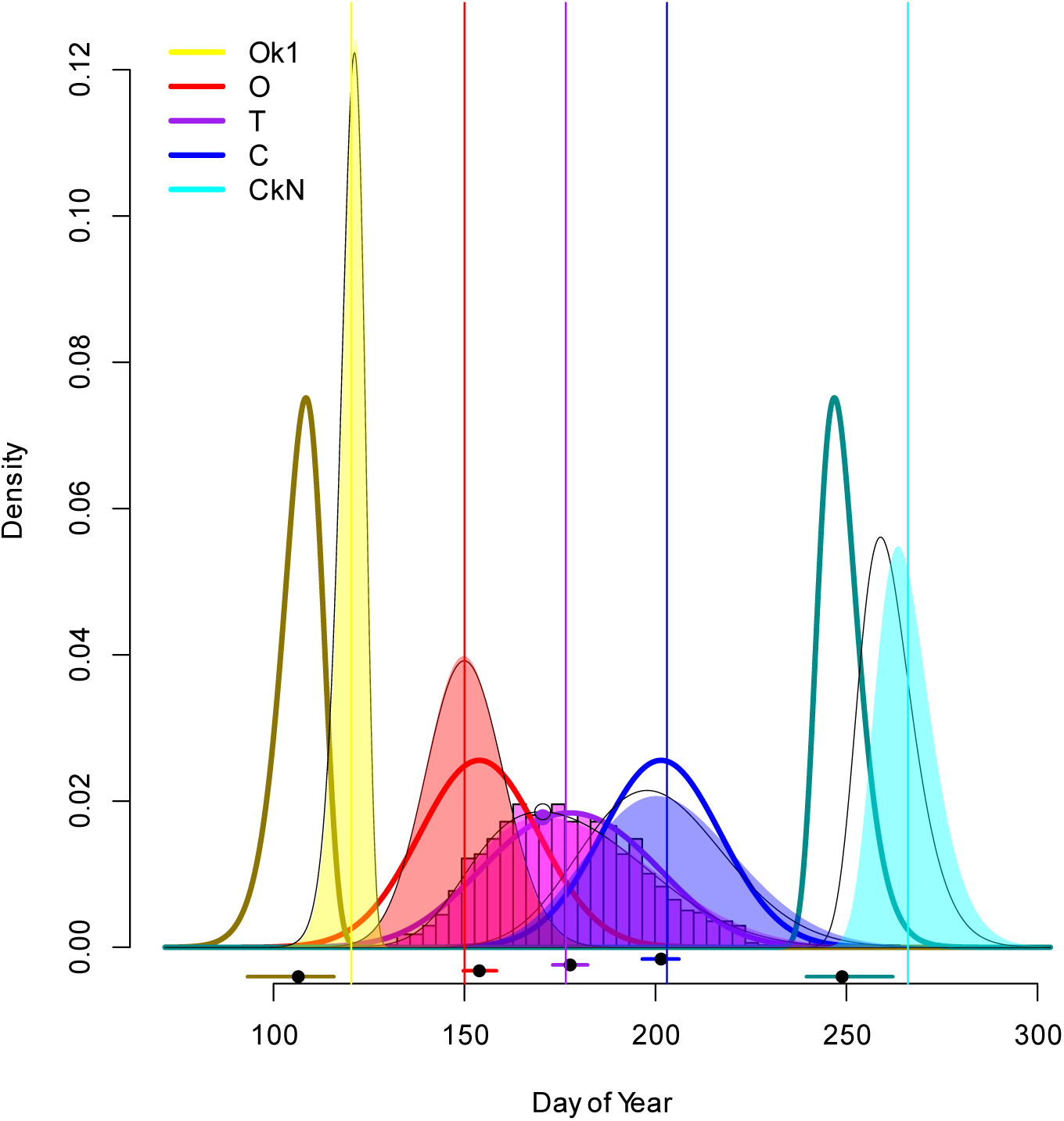
Pathological case of model mis-specification. Shaded regions represent theoretical probability distributions based on the BB model, using the same parameters as in in Fig. 3. Thick, solid, colored curves show probability density functions (PDFs) based on posterior mean parameter estimates inferred using Bayesian GP, which employed a mis-specified GP model to BB generated data. Thin black lines represent maximum a posteriori (MAP) estimates obtained via numerical optimization under the BB model, the true data-generating model. Closed and open circles denote the true and MAP-estimated values, respectively, for the peak of the phenophase. Horizontal intervals beneath the distributions indicate Bayesian 95% credible intervals (CIs) for the means of the respective distributions. Black dots in these intervals mark the posterior means, whereas vertical colored lines indicate the true parameter values. The true values fall within the 95% CIs for the onset *O*, observed-time *T*, and cessation *C* distributions despite model mis-match.

**Fig. 5.**
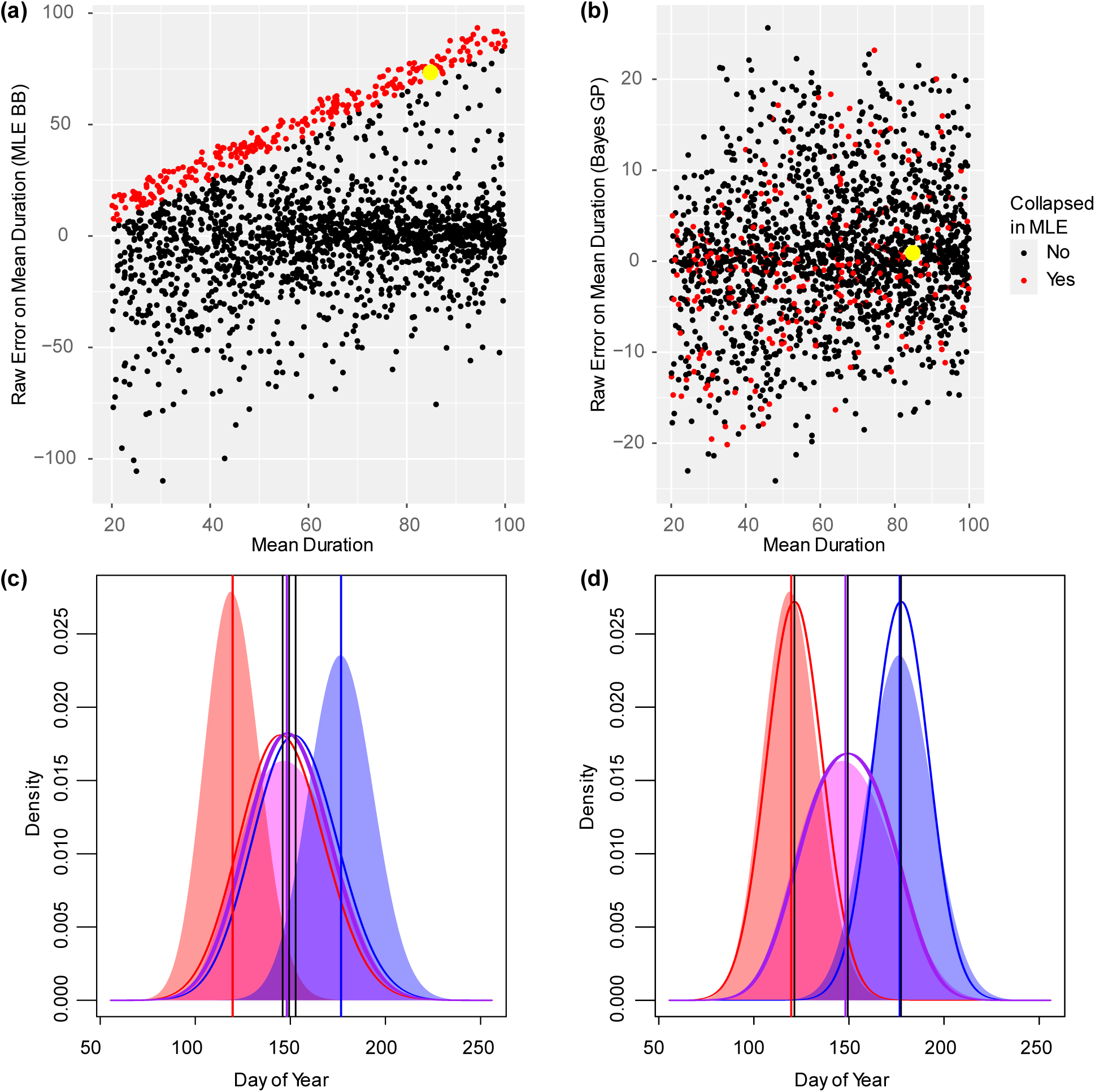
Model identifiability and the role of prior distributions. (a), (b) Each point represents a different simulation replicate generated using randomly selected parameter values as described in the text and Table S1. (a) Raw error in estimating µ_D_ using maximum likelihood estimation (MLE). Multiple parameterizations can closely fit the observed data, but some near-optimal solutions yield implausibly small estimates of µ_D_ (shown as red dots), resulting in “collapsed” durations. (b) The same simulation replicates, but with parameters estimated via Bayesian GP. The issue of collapsed durations is resolved, and absolute error is reduced on average compared to MLE. (c) True distributions (solid shaded regions) and MLE-based fits (curved colored lines) for the simulation corresponding to the yellow dot in panels (a) and (b), illustrating the collapsed duration issue. Data underpinning the yellow dot were simulated under the mis-matched BB model. Solid, colored vertical lines indicate true parameter values; thin black lines indicate MLE estimates. (d) True distributions (solid shaded regions) and mean posterior –based fits (curved colored lines) for the same simulation (yellow dot). The collapsed duration issue is eliminated.

There was also a significant difference for peak phenophase. The peak phenophase under the GP is in the middle of the observed-time *T* distribution because it is symmetrical. However, when the observed-time *T* distribution is not symmetrical, the peak will not be in the middle, and Bayesian GP inferences will be incorrect.

In cases where the observed-time *T* distribution is skewed and researchers are particularly interested in peak and extreme events, there are alternative approaches. One alternative is to estimate the parameter values that maximize the posterior distribution for the BB model. The thin black lines in Fig. 4 illustrate results of this approach. We were unable to implement the BB model as a Stan probability model due to numerical stability issues and high divergence rates, so this approach only provides point estimates, it does not assess parameter uncertainty, and covariates cannot be included in our implementation. MLE is not recommended for reasons presented in the next section.

### Model Identifiability and Priors

Maximum likelihood estimation (MLE) provided relatively poor estimates due in large part to model identifiability issues (Fig. 5). One particularly pernicious issue was that MLE often resulted in estimates of durations μ_*D*_that were much smaller than true values (i.e., ‘collapsed’; Fig. 5a). Nevertheless, the inferred distribution (purple curve in Fig. 5c and d) matched the true observed-time *T* distribution (shaded purple in Fig. 5c and d) relatively well, thereby illustrating model non-identifiability. Indeed, in some cases, the likelihood for the ‘collapsed’ model was as high as, or even higher than, the likelihood at the true parameter values. However, this difference in likelihood was smaller than the contribution that the prior makes to the posterior probability. Thus, incorporating an informative prior resolves the issue (Fig. 5b, d). An informative prior helps constrain the inference, discouraging estimates that deviate substantially from the true parameters and mitigating identifiability issues.

The prior distribution influences parameter estimation error (Fig. 6). The difference between the mean of the prior distribution and the true parameter value (i.e., hyperparameter bias) is directly correlated with error rate. For example, hyperparameter bias is linearly correlated with error of μ_*D*_ estimates (Fig. 6, β=0.54, *t*(1639) = 23.5, *p*-value < 0.001). Despite some simulation replicates exhibiting hyperparameter biases of up to 10 days, approximately 92% of replicates yielded estimates within the 95% Bayesian credible interval. Incorporating data via the likelihood function in the Bayesian analysis thereby reduces error generated by biased hyperparameters by nearly one half on average, marginalized across parameter, hyperparameter, and sample size variation, as reflected by the regression slope of approximately ½ in Fig 6. Theoretical results (Edwards *et al*., 1963) indicate that with informative, but not overly peaked priors that are non-zero for the range of feasible parameter values, the signal from the prior will eventually be overcome by the weight of empirical data (the error slope will go to zero). When hyperparameter bias is zero, the mean estimation error is not significantly different from zero (estimate of intercept = –0.009849, *t* = –0.074, *p*-value = 0.941), indicating that the Bayesian GP inference procedure is unbiased when priors themselves are unbiased.

**Fig. 6.**
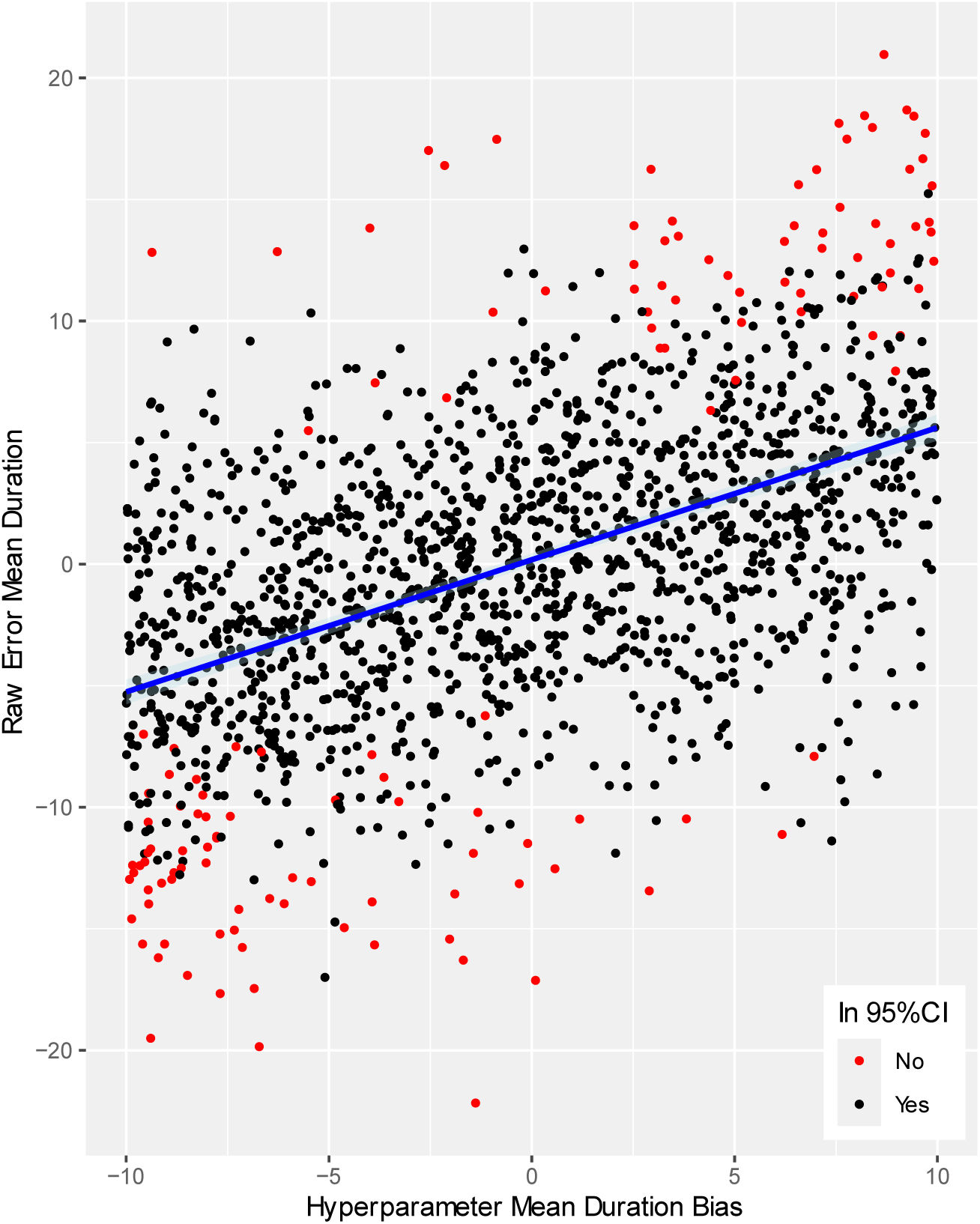
Influence of hyperparameter bias on parameter estimates. Each point represents a simulation replicate in which both parameters and hyperparameters were randomly selected. The plot shows raw error (in days) for estimates of μ_D_ as a function of bias (in days) in the hyperparameter mean for μ_D_. The blue line is the linear regression fit, with a narrow shaded region showing the 95% frequentist confidence interval for the mean. Red points indicate simulation replicates for which the estimated value of μ_D_ falls outside the 95% Bayesian credible interval.

These results suggest that the prior distribution is a key component of the success of the Bayesian GP methodology. In later sections, we illustrate an approach that provides fairly accurate hyperparameter estimates in an automated fashion, making it feasible to implement these methods without requiring substantial prior experience.

### Comparison to Pearse Estimator (PE) of Phenological Extremes

Bayesian GP estimates for phenological extremes converged to zero mean raw error with sample sizes of around 60 or higher (Fig. 7). As was previously recognized by Cooke (Cooke, 1979, 1980) for order statistics, estimates were biased and undershot true values for small sample sizes, but estimates based on sample sizes of at least 60 individuals appear to be unaffected by bias; the dotted lines representing the estimates converged to the true parameter values (Fig. 7a).

Our conceptual framework recognizes onset and cessation distributions as separate from distributions of phenological extremes. The distributions of phenological extremes are population level distributions: multiple different populations are required to generate a histogram based on empirical data. In contrast, phenological metrics of individuals from a single population are sufficient to generate a histogram for onset and cessation distributions. Fig. 1 illustrates this idea with the single black lines at the periphery of the onset and cessation histograms.

Our simulations suggest that the Pearse et al. estimator (PE) is insensitive to considerations of population size. Estimates were largely constant across sample sizes and population sizes and did not reflect the true first onset value (Fig. 7b). These results indicate that when using the PE method, the average estimate of first onset remains the same regardless of the population size. For example, drawing a random sample of 10 individuals from a population of 1,000,000 yields, on average, the same PE estimate as drawing 10 individuals from a population of 100, as illustrated in Fig. 7b. In all scenarios investigated, PE consistently exhibited bias. At small sample sizes, confidence intervals were wide enough to include the true value, irrespective of population size. Pearse et al. applied this method to small datasets; in their Fig. 3, 5 of the 7 species were represented by samples of 10 or fewer individuals (none exceeded 60). This suggests that their confidence intervals may have captured the true value primarily due to the high uncertainty associated with small sample sizes, rather than the accuracy of the estimator.

A key result is that analyses of phenological extremes are more likely to obtain accurate estimates of phenological extremes when population sizes are accounted for. Phenological extremes are functions of population size, as indicated by equations [8] and [9]. The connections between population size and estimation of phenological extremes has been recognized for quite some time (Miller-Rushing *et al*., 2008; Moussus *et al*., 2010). Primack et al. (Primack *et al*., 2023) illustrate this relationship between population size and phenological extremes in their Fig. 2.

Estimating population size may not be especially problematic. Equation [11] indicates that phenological extremes are well-approximated by a function of the inverse CDF Φ^−1^(*p*) of the normal distribution. Furthermore, Φ^−1^(*p*) is asymptotically bound by 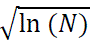 (Blair *et al*., 1976). The square root of the logarithm of a number increases slowly as *N* increases, so phenological extremes are expected to be quite similar for a broad range of larger population sizes. This idea is also depicted in Fig. 8 by comparing the distance between the true first onset for a population of *N=100* (top black horizontal line) to that of the population of *N=*10,000 (middle line), which is *larger* than the distance between the true value for the population of *N=*10,000 (middle line) to that of the population of *N*=1,000,000 (bottom line). The difference in estimates for *N=*10,000 and *N*=1,000,000 is only around 10 days under this specific set of population parameters. This result is helpful, because it suggests that estimates of population sizes can have a large error but estimates of phenological extremes will nonetheless be fairly accurate using Bayesian GP.

**Fig. 8.**
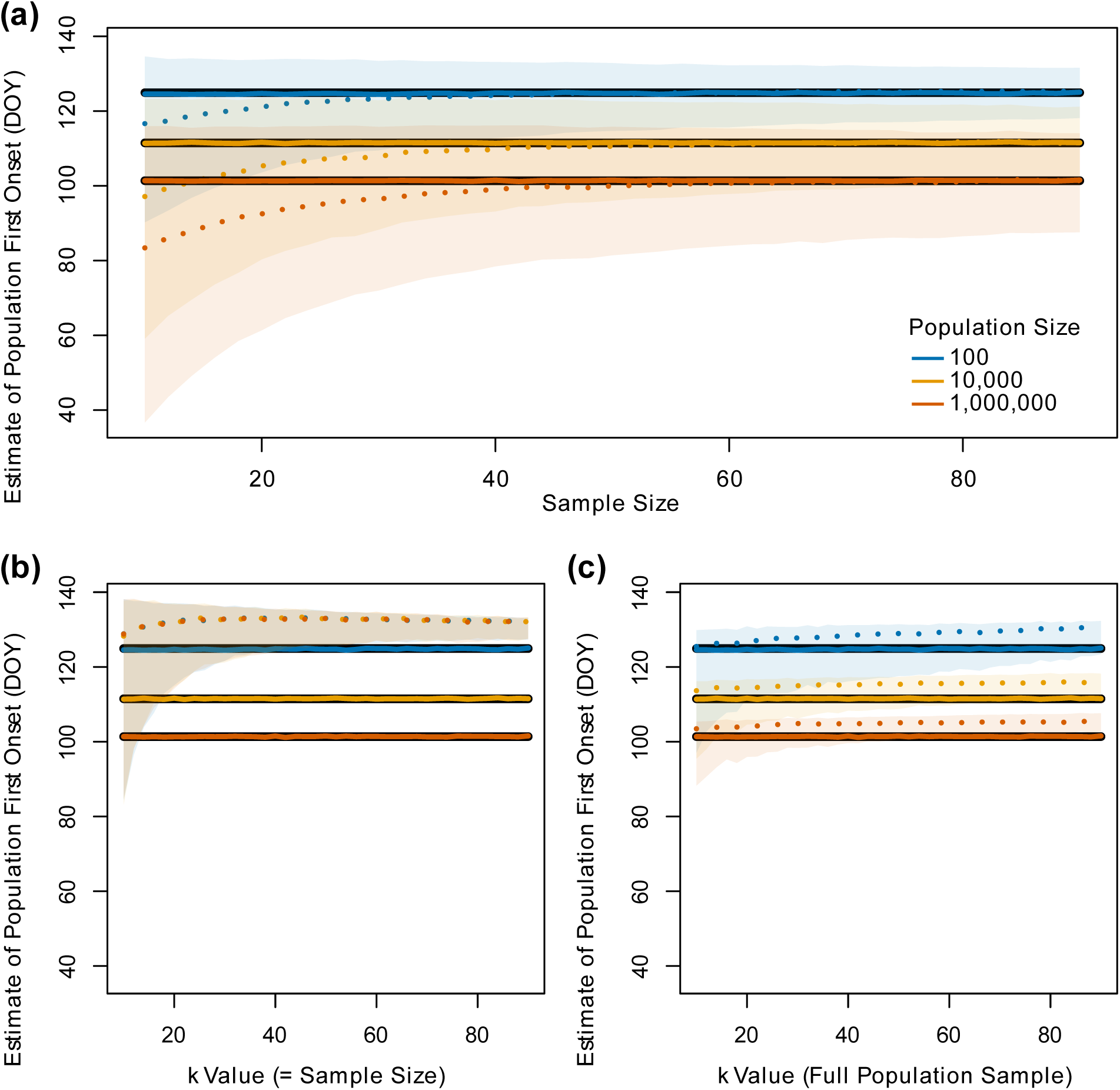
Bayesian GP inferences of phenological extremes compared to the Pearse et al. (2017; PE) estimator. Estimation raw error for the mean first onset μ_*O*1_ was compared between the PE method and Bayesian GP across varying sample sizes (*x*-axis) and simulated population sizes (*N* = 100, 10,000, and 1,000,000). Each combination of population size and sample size was simulated 1,000 times. Individuals were randomly subsampled according the sample from the simulated population of 1000 individuals and used to make inferences. Theoretical estimates μ_*O*1_ based on Equation [8] are shown as black solid lines. The earliest onset time in the simulated population was averaged across the 1,000 simulation replicates per scenario and plotted as solid, colored lines (closely aligned with the theoretical expectations). Simulation parameters were set to µ_O_ = 150 days, µ_D_ = 30 days, and σ = 10 days. Dotted lines show mean estimates, and shaded regions represent 95% credible intervals (a) or 95% confidence intervals (b, c). (a) Bayesian GP results. For each replicate, the mean of the posterior distribution for μ_*O*1_ was used as the point estimate represented by the dotted line, and the 95% credible interval was calculated by averaging lower and upper CI bounds across simulation replicates. (b) PE estimates using sampled data. The `weib.limit` function from Pearse et al. was applied to the sample of observed times. The *k* parameter (number of observations used for estimation) was set equal to the sample size, and α = 0.05 defined the confidence interval. (c) PE estimates using full population data. The *k* value varied according to the x-axis values, but the full simulated population was provided as the sample.

The extreme value theorem (EVT), also known as the Fisher-Tippett-Gnedenko theorem (Fisher & Tippett, 1928; Gnedenko, 1943), is a limit theorem in probability that describes the asymptotic behavior of the maximum (or minimum) value of a large population. Provided that a non-degenerate limit exists, under appropriate scaling and translation of the extreme value, the distribution of phenological extremes converges to one of three possible families of distributions: Weibull, Fréchet, or Gumbel. Convergence to the Weibull type occurs when the cessation or onset distribution is bounded and has light (exponentially-decaying) tails. These conditions are particularly relevant for phenological extremes, which are often constrained by hard biological or environmental limits (e.g., end of a growing season when freezing occurs).

Beta, uniform, and truncated normal are examples of such distributions that are bounded and have light tails. As such, we expect phenological extremes to be well-modeled by the Weibull, suggesting that distributions in equations [8] and [9] converge to the Weibull for appropriate underlying onset and cessation PDFs. However, for the standard normal distribution that has unbounded support, EVT predicts convergence to a Gumbel-type extreme value distribution, not Weibull. However, from a biological standpoint, when environmental factors (such as winter freezing) impose hard boundaries on phenological behavior, the Weibull distribution is arguably the most appropriate model for the distribution of extremes.

Pearse et al. (Pearse *et al*., 2017) employed an unusual case in their simulations. They used a Dirac delta distribution for the onset and a Dirac delta distribution for the cessation. A Dirac delta distribution has zero variance (all probability mass is concentrated at a single point), meaning that the Dirac delta distribution is degenerate – EVT doesn’t apply! Under their model, PE frequently produces estimates that are outside the range of possible values that fall between the hard onset threshold and the hard cessation threshold. In other words, their procedure can produce estimates than have zero probability under their model.

This degenerate case provides an interesting example, because it is one of the few cases when the distribution of observed times *T* is a standard distribution: the uniform distribution. We are unaware of empirical phenological distributions that match these assumptions.

Pearse et al. (Pearse *et al*., 2017) made seminal contributions by linking phenological extremes to the Weibull distribution, insights that later informed the development of the *phenesse* package (Belitz *et al*., 2020). Belitz et al. (Belitz *et al*., 2020) used Cooke’s (Cooke, 1980) result that the tails of distributions derived from sparsely sampled phenological data often resemble Weibull distributions. Consequently, *phenesse* fits a Weibull distribution to the observed-time *T* distribution and provides quantile estimates, using a bootstrapping procedure to reduce estimation bias. While quantile-based approaches are useful for estimating phenological thresholds (e.g., the date by which a certain percentage of events have occurred), we advocate for Bayesian methods that provide full posterior distributions for observations (posterior predictive probabilities) and parameter estimates, avoiding the need for potentially arbitrary cutoff values.

### Analysis of the Phenology Paradox using Bayesian Inference, Quantile Regression, and Standard Linear Regression

Like Belitz et al. (Belitz *et al*., 2020), Park et al. (Park *et al*., 2025) also explore the use of quantiles as estimators for phenological events. They recognize that their cutoffs are “arbitrary” (their wording), and use the 10% quantile for the onset, the 50% quantile for the peak, and the 90% quantile for the cessation in their quantile regression (QR) approach. Like us, they acknowledge distinct onset, observed, and cessation distributions, and they use an approach to simulate data that is similar to ours and what Hearn (Hearn, 2022) presented previously. As one of the few methods that distinguishes among these phenological distributions, we chose to compare accuracy of our method to theirs. As an onset distribution-aware procedure, we were particularly curious to explore if their method accurately detects the signs of slopes for each of the six Scenarios (Fig. 2).

QR performs comparably to Bayesian GP to detect slopes for linear models having one covariate (Fig. 8). Because it provides close approximations to the slopes, we used QR estimates as hyperparameter values. For all Scenarios and all slope types, our method has lower absolute error on average. By setting priors at QR estimates, the Bayesian GP method further refines estimates. When SR is used to estimate the onset slope, it fails in almost every simulation replicate to correctly identify the sign of the onset model slope for the paradoxical Scenarios 3 and 6 (2 out of 1000 simulation replicates). Both QR and Bayesian GP resulted in lower error on slope parameters than SR.

Although quantile regression (QR) accurately estimates slope coefficients, it was less effective at estimating the anchors of linear models. In linear regression, the anchor corresponds to the marginal mean of the response and reflects the vertical position—or shift—of the regression line. Since the mean regression line passes through the point defined by the mean response and mean covariate, accurately estimating the anchor is essential for capturing the overall level along the response variable axis of the relationship. As the mean duration μ_*D*_increases, accuracy of the estimate for μ_*O*_ decreases, on average, when QR is used (Fig. 9; β = 0.11, *p*-value < 0.001), indicating that extended phenophase durations lead to less precise quantile estimates. Accuracy of the estimate for μ_*O*_ is not correlated with μ_*D*_ under Bayesian GP (β = –0.0017, *p*-value = 0.55). Bayesian GP achieved significantly lower error (mean = 1.49 days) compared to QR (mean = 4.3 days; t = 31.201, *df* = 1502, *p*-value < 2.2 × 10⁻¹⁶). Bayesian GP yielded lower onset anchor error than QR in ∼82.6% of replicates. Both methods exhibited significant correlations between error on onset anchor and sigma (σ), but in opposite directions (GP: *t* = 7.329, p = 4.78 × 10⁻¹³; QR: *t* = –19.695, p < 2 × 10⁻¹⁶). Errors in onset slope estimates were also significantly associated with both duration (QR: F₁,₉₉₈ = 17.22, p = 3.62 × 10⁻⁵; QR: F₁,₉₉₈ = 13.32, p = 2.76 × 10⁻⁴) and σ (GP: F₁,₉₉₈ = 30.2, p = 4.94 × 10⁻⁸; QR: F₁,₉₉₈ = 24.92, p = 7.05 × 10⁻⁷). However, the estimated regression coefficients (in days per unit covariate) for both duration (GP: 1.309 × 10⁻⁴; QR: 1.668 × 10⁻⁴) and σ (GP: 9.067 × 10⁻⁴; QR: 1.204 × 10⁻³) suggest limited practical effect despite statistical significance.

**Fig. 9.**
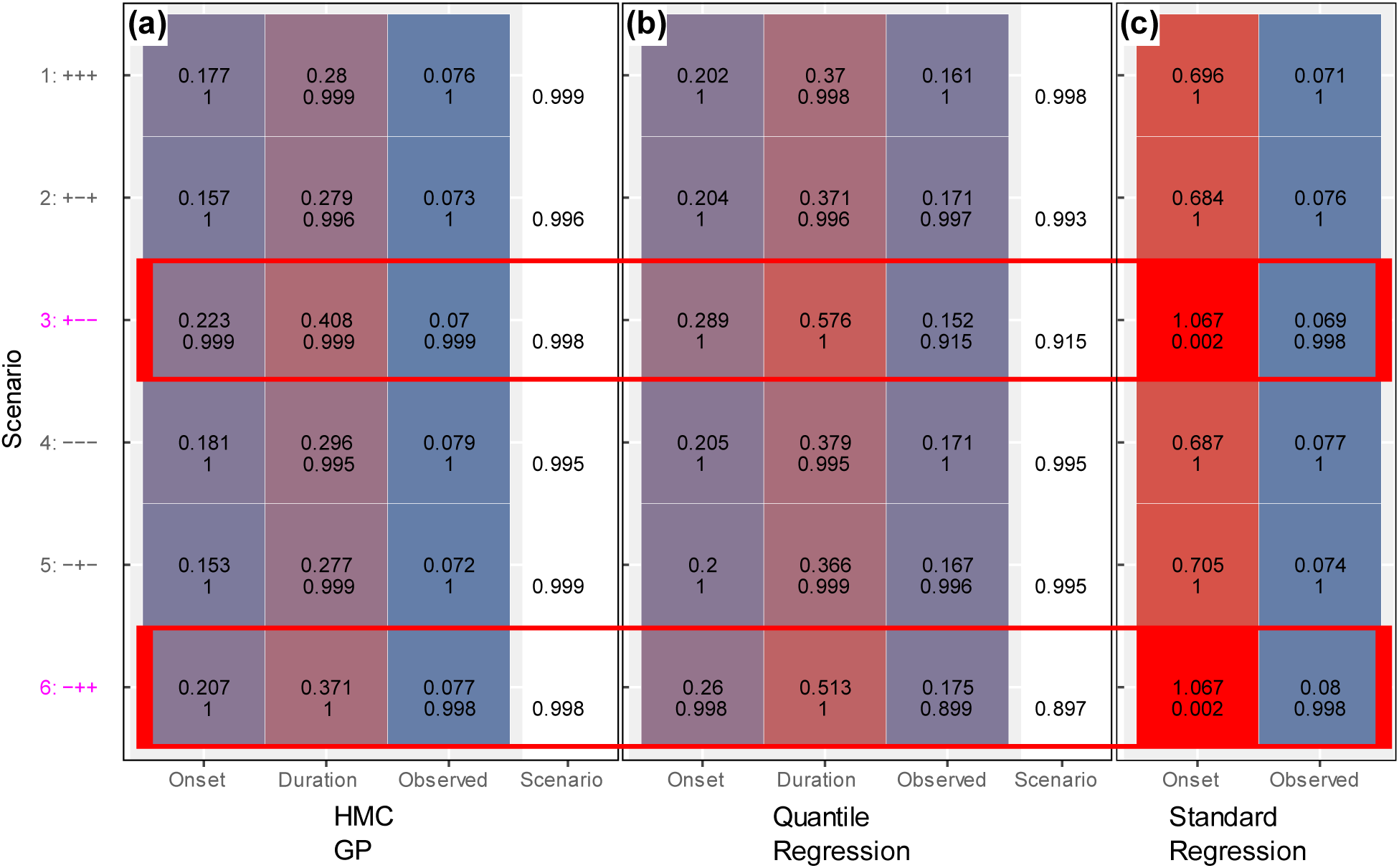
Evaluation of methods for detecting phenological Scenarios. For each of six Scenarios (numbered as in Fig. 2, with 3 and 6 classified as “paradoxical”), 1,000 simulation replicates were conducted (detailed in Table S1). Each subpanel (a–c) summarizes mean estimation accuracy and classification performance for different inference methods. The Onset, Duration, and Observed columns report absolute error (top number) and proportion of replicates with correctly inferred slope sign (bottom number). The Scenario column shows the proportion of replicates in which the method correctly identified the directional signs (±) for onset, duration, and observed-time that occur for the Scenario. (a) Bayesian GP using HMC sampling. (b) Quantile regression (QR). The 10th and 90th percentiles were used to model onset and cessation, respectively; the observed-time model was the 50th percentile. Duration slopes were derived from onset and cessation models by taking their difference. (c) Standard linear regression (SR) on observed values only. Slope estimates from the observed model were used as proxies for onset slopes as has been done in the literature. Bayesian GP yielded the lowest absolute error across all categories. Paradoxical scenarios had the highest onset errors, particularly under SR, for which the correct onset slope sign was identified in only 0.2% of those replicates.

Because quantile regression (QR) is highly automated, it offers a convenient method for generating initial hyperparameter estimates without requiring prior knowledge. For sufficiently large datasets (e.g., with more than 100 records), we recommend partitioning the data: using 30% to estimate prior hyperparameters via QR, and the remaining 70% for the full Bayesian Gaussian Process (GP) analysis. While MAP optimization offers a potential method for estimating anchor values of the hyperparameters, this approach was not explored in the present study.

### Analyses of Empirical Datasets

Our primary goal in analyzing the empirical datasets was to assess how often changes in duration contribute to the mismatch between slope estimates from SR methods and the true slopes of the onset model. Above, we showed that both changes in onset and changes in duration influence the observed pattern of specimen collection times, and SR cannot tease apart the relative contributions of these two sources of variation. We also showed that when durations do not change, SR estimates of parameters are valid estimates of onset model parameters. So, empirically, if there are no changes in duration, SR estimates are sufficient. By explicitly incorporating separate latent variable models of the onset and duration, our approach can tease apart the relative contributions of onset and duration to the observed pattern of specimen collection DOY.

In total, we used 5,363 specimens for the empirical analyses, starting with 12,125 specimens, many of which lacked images to assess phenological state. 680 specimens were removed due to incomplete records, duplicated entries, or identified outliers. While removing outliers may eliminate valid observations and potentially bias results by underestimating true variability, a detailed investigation of these outliers is beyond the scope of this study. Summary statistics for the empirical data are available in Table S2.

To assess the posterior probability of a change in duration associated with a covariate, we calculated the proportion of posterior draws in which the slope was negative (Table S3, column ‘posterior.prob.neg.slope’). This represents the probability that duration decreases as the covariate increases, marginalized over other factors. By the complement rule, one minus this probability gives the probability that duration increases with the covariate. In our dataset, every species exhibited a change in slope with a posterior probability of at least 0.95 for at least one covariate, providing strong evidence that duration is influenced by environmental factors such as elevation, latitude, spring temperature, or annual temperature. Consequently, the slopes from the onset model differed from those of the observed-time model. These patterns are illustrated in Fig. 13.

**Fig. 10.**
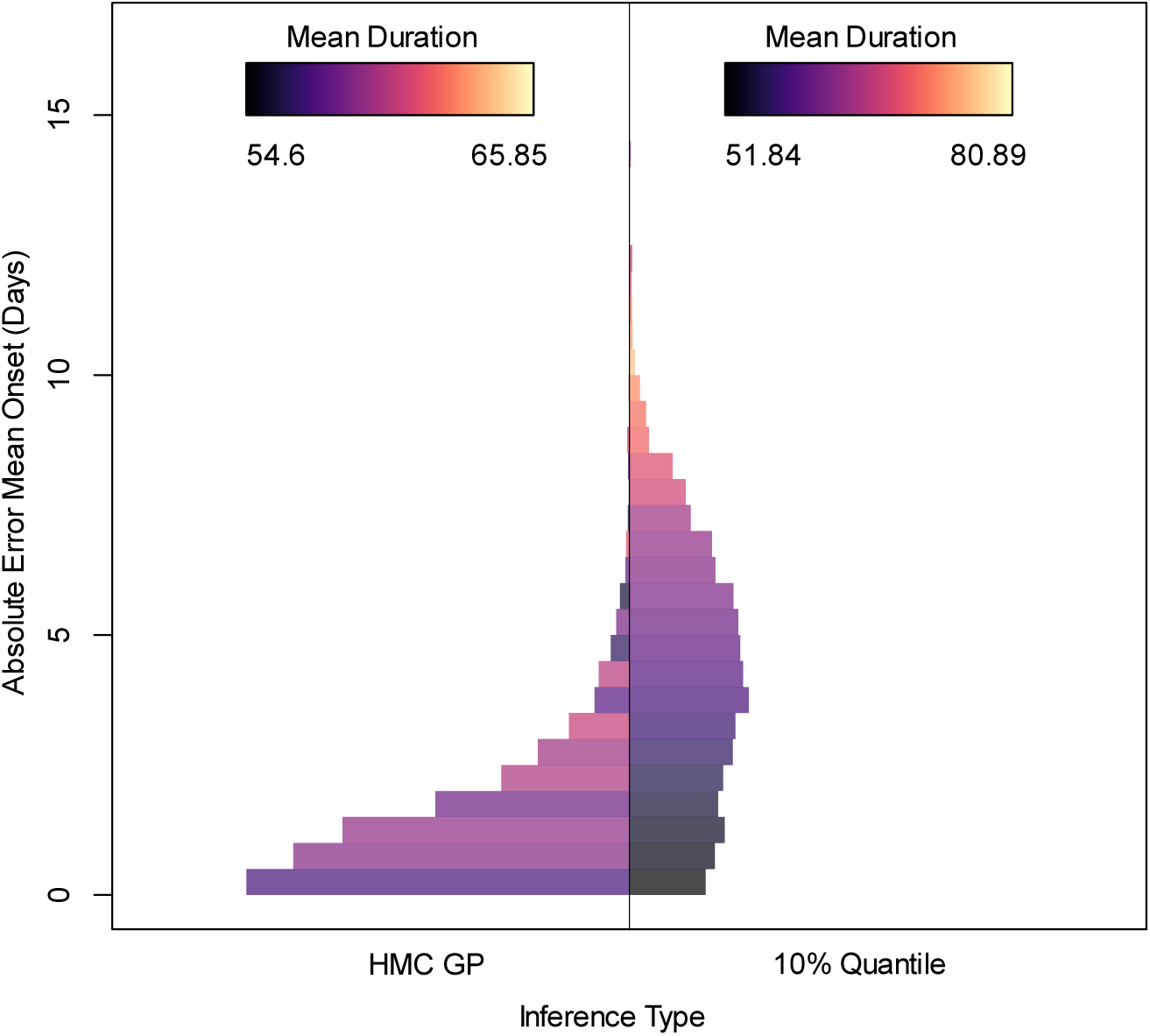
Comparison of absolute error in estimates of mean onset between Bayesian GP and quantile regression (QR), highlighting the influence of duration. The histograms show the absolute error for the onset model anchor across 1,000 simulation replicates, marginalized over randomly sampled model parameters, hyperparameters, and sample sizes. Histogram bars are colored by the mean of the duration anchor for the replicates summarized within each bin; the bar length (along the *x*-axis) represents the frequency of simulation replicates falling into the error range depicted on the *y*-axis. Bars on the left represent errors from Bayesian GP posterior mean estimates, and those on the right represent errors from QR estimates. The correlation between error and duration is evident by the coloring of bars for QR estimates.

**Fig. 12.**
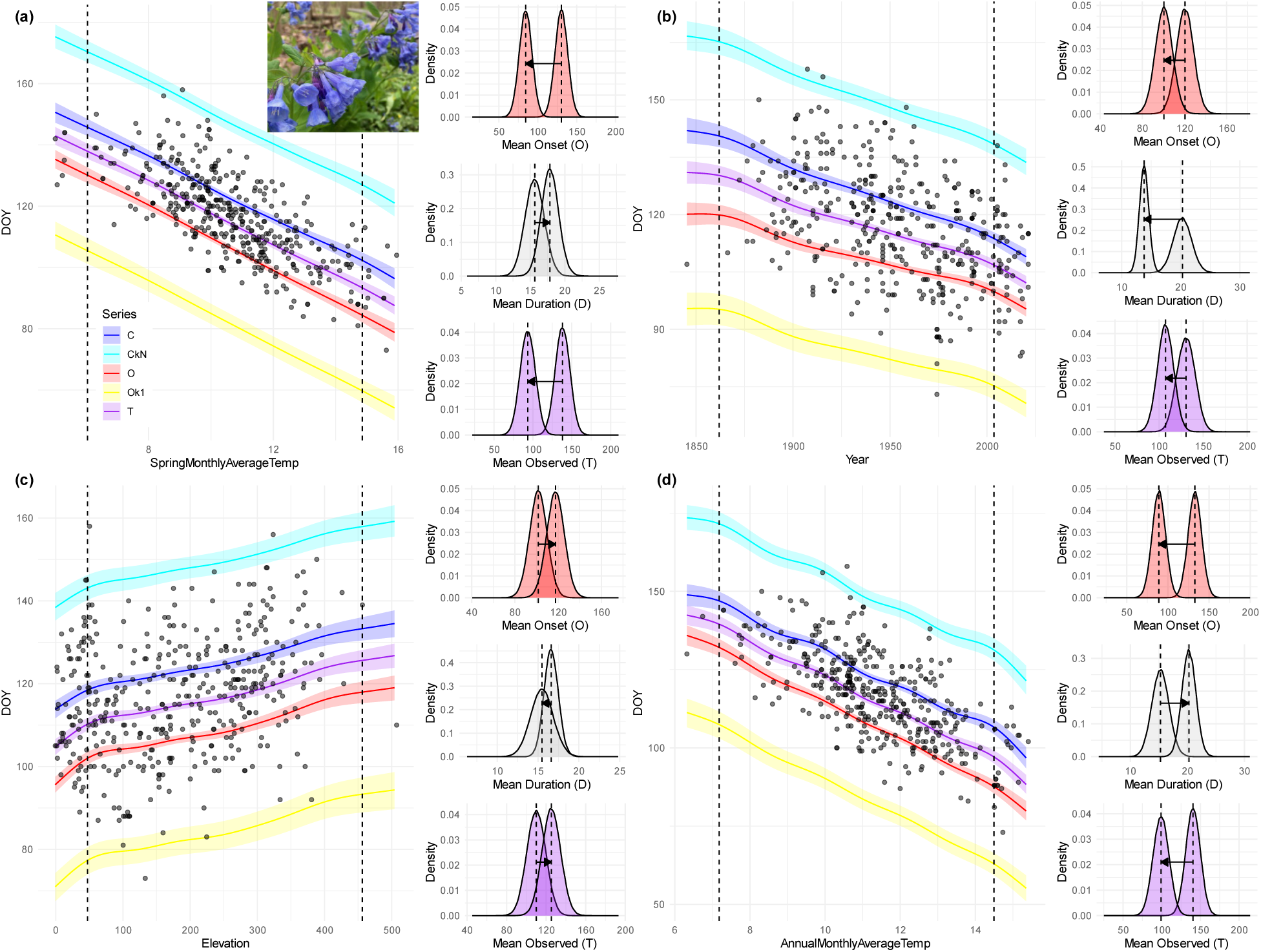
Posterior predictive plots for *Mertensia virginica*. Each panel (a–d) shows empirical and model-based variation in phenology across one of four covariates. The left side of each panel is a scatterplot of observed collection day of year (DOY) against: (a) spring monthly average temperature,(b) year, (c) elevation, and (d) annual monthly average temperature. Color-coding is the same as previous figures. Shaded ribbons represent 95% credible intervals for the mean predicted DOY. As expected, data-rich regions of covariate space result in narrower 95% CI’s whereas data-poor regions result in wider intervals. Vertical dotted lines indicate two covariate values at which the posterior predictive distributions (PPDs) shown on the right side of each panel were made. These PPDs are marginalized over all other covariates and model uncertainty. For each covariate slice, three distributions are shown (top to bottom): predicted day of onset, predicted duration, and predicted day of cessation. Arrows illustrate the directional shift from the lower to the higher covariate value. Scenario 5 (-+-) is evident in (a) and (d), scenario 4 (---) is present in (b), and scenario 2 (+-+) is observed in (c). Photo by Geoffrey A. Landis, https://creativecommons.org/licenses/by-sa/4.0/

**Fig. 13.**
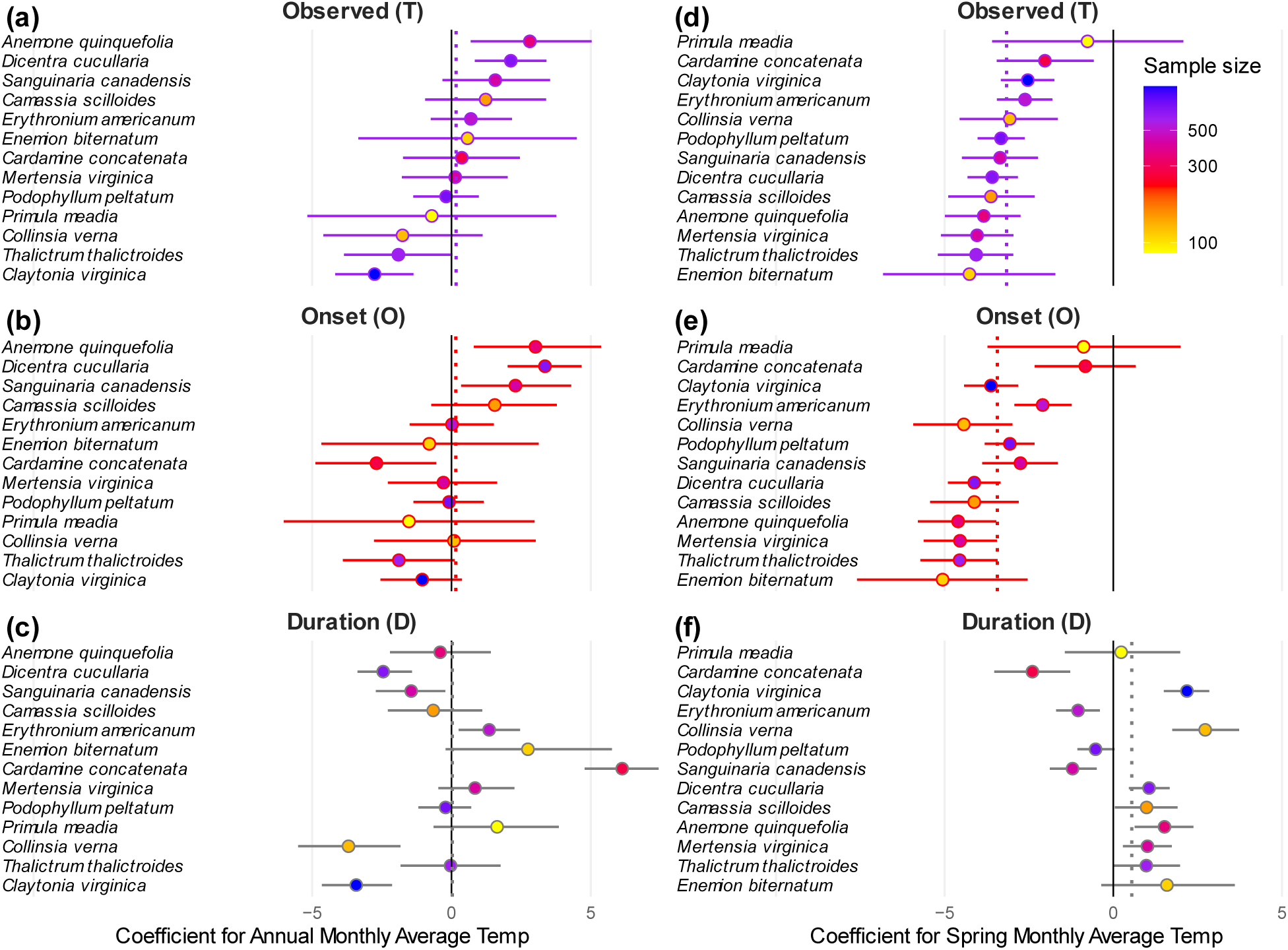
Summary of empirical analyses. Panels show posterior estimates of slope coefficients for models using either Annual Monthly Average Temperature (left column: a–c) or Spring Monthly Average Temperature (right column: d–f) as covariates, with subpanels for (a, d) observed time, (b, e) onset, and (c, f) duration. Species are ordered from highest to lowest slope for the observed-time model. Each dot represents the posterior mean for a species-specific slope coefficient; horizontal bars indicate 95% credible intervals. Dot colors indicate sample size, as shown in the legend.

**Fig. 14.**
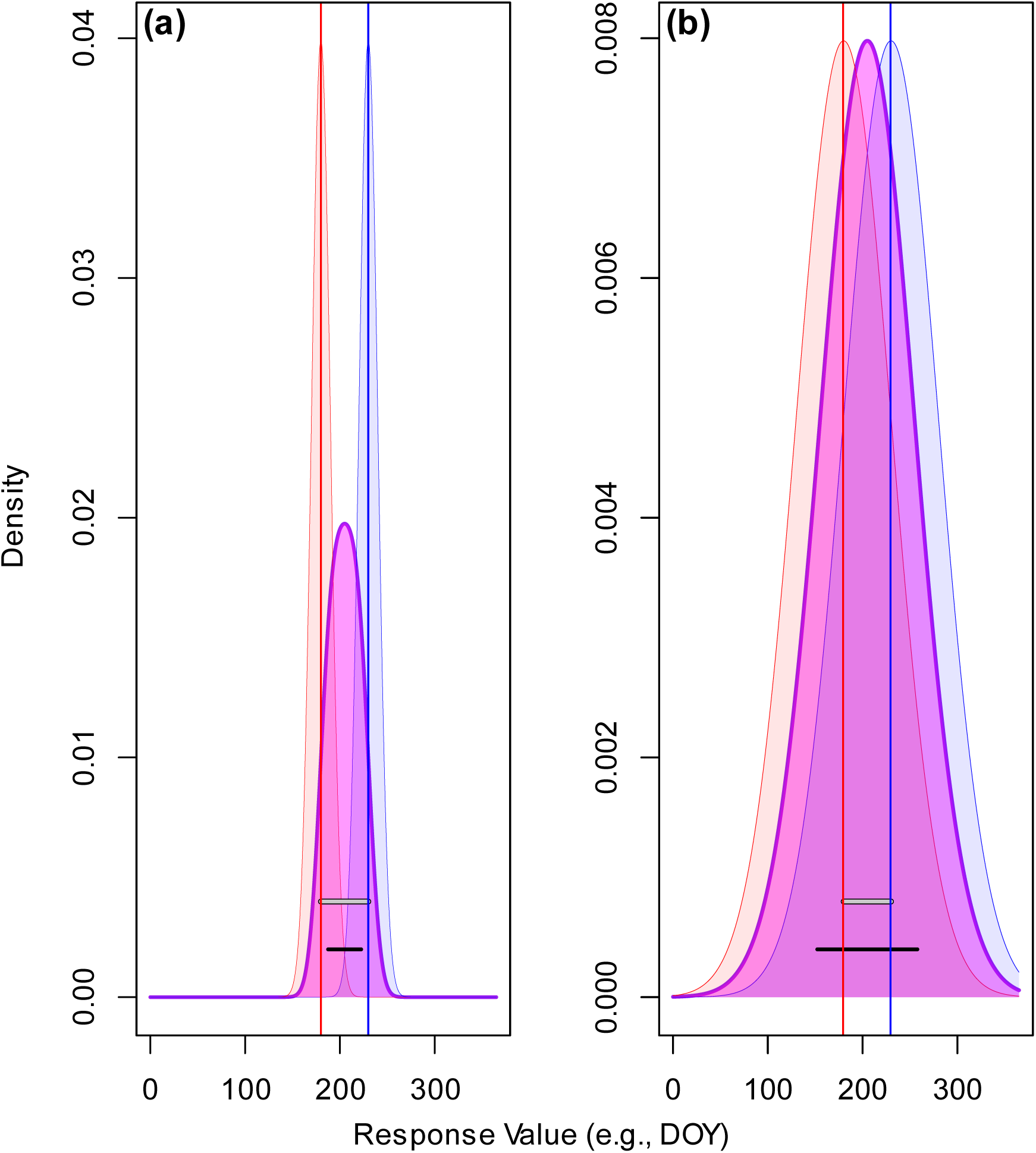
Relationship between σ and duration: why variance in the observed-time distribution is not a proxy for phenophase duration. The width of the observed-time distribution results from a combination of variation in onset times (σ) and duration. This figure illustrates two scenarios in (a) and (b) where the mean duration remains constant (gray horizontal bars), but the variation in onset and cessation differ between these two scenarios, resulting in markedly different variation in observed times (black horizontal bars). (a) When onset and cessation times vary little across individuals, observed times can be tightly clustered. (b) When onset and cessation times exhibit greater variability, the observed-time distribution is correspondingly wider, even though the mean duration has not changed in this example. Thus, variance or standard deviation of observed times cannot be used as a reliable surrogate for phenophase duration, since it conflates multiple sources of temporal variation. Colors follow those used in previous figures.

We did not observe any species that exhibited paradoxical scenarios, although *Cardamine concatenata* is a candidate for Scenario 6. However, its 95% CI for the observed-time slope straddles zero. Our analysis was limited to only 13 species, so it is difficult to assess the frequency of the paradoxical situations. In the absence of paradoxical behavior, slopes from the onset model are expected to correlate with those from the observed-time model, though they are not necessarily identical. Indeed, we found a positive correlation between onset model slopes and observed-time slopes. For example, the onset slopes corresponding to Spring Monthly Average Temperature were positively correlated between SR estimates and Bayesian GP (F-statistic = 36.39 on 1 and 11 *df.* Adjusted R^2^: 0.7468, *p*-value = 8.525e-05) and correlated for Annual Monthly Average Temperature as well (F-statistic = 12.18 on 1 and 11 *df*. Adjusted R^2^: 0.4824, *p*-value = 0.005056). Because the regression R^2^ values are not equal to one, substituting observed-time model coefficients as proxies for onset model slopes unnecessarily introduces error. Thus, assessment of phenological sensitivities is more accurately assessed using Bayesian GP.

One way to assess the quality of Bayesian GP estimates is to compare is to compare them to those estimated using standard regression (SR). As shown above, β_***T***_ estimated using SR should agree with 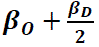 estimated from Bayesian GP. Agreement between these estimates would indicate consistency and reliability of the Bayesian GP approach. In all cases, we found that the SR estimates fell within the 95% credible interval (CI) of the Bayesian estimates for β_***T***_(Table S3, column ‘in.95CI’). Initially, the Bayesian GP 95% CIs for β_***T***_ for *Primula meadia* and *Enemion biternatum* did not fully cover the SR estimates. These two species had the smallest sample sizes in our dataset (Fig. 13; Table S2), and Stan produced inconsistent results across different runs. To improve sampling and convergence, we doubled the warmup and sampling phases to 2,000 iterations each. Following this adjustment (reflected by increased effective sample sizes (ESS) for these species in Table S2) the Bayesian GP CIs overlapped with the SR estimates. Fig. 13 highlights that 95% CI lengths decrease on average as sample sizes increase, reflecting greater precision as more data are added. Linear regression confirms a statistically significant negative relationship (α = 0.01) between sample size and CI interval length across all combinations of model type (onset, observed, duration) and covariate (annual and spring temperature depicted in the figure) combinations.

Sampling diagnostics for all species were within acceptable ranges. The percentage of divergent transitions was zero, E-BFMI values all exceeded 0.3, effective sample sizes (ESS) were in the high hundreds or greater, and all R-hat values were close to 1. ESS and R-hat values for individual parameters are provided in Table S3. These results were not achieved immediately. Initial model implementations exhibited unacceptably high divergence rates. Reparametrizing linear models in terms of an anchor rather than an intercept helped mitigate some issues, but the most effective strategy was the specification of priors for both population-level and individual-level parameters, thereby implementing a hierarchical Bayesian model. We believe the improved diagnostics resulted from the regularizing influence of these priors, which would smooth the posterior distribution and reduce abrupt changes in parameter values. This, in turn, likely improved the performance of Stan’s automatic differentiation algorithms. While we did not explore this effect further, the result may offer general guidance for others developing phenology models in Stan.

Based on the results of the above analyses of phenology paradoxes (*Analysis of the Phenology Paradox using Bayesian Inference, Quantile Regression, and Standard Linear Regression*), we used QR to specify hyperparameter values in our analyses of empirical data. Although there are multiple studies of spring ephemerals that highlight phenological processes that we could use for priors (e.g. (Oldham, 1990; Abu-Asab *et al*., 2001; Badeck *et al*., 2004; Primack *et al*., 2004a; Calinger *et al*., 2013; Pearson, 2019c; Petrauski *et al*., 2019; Alecrim *et al*., 2023; Miller *et al*., 2023; Faidiga *et al*., 2023; Watson & Vuorisalo, 2024; Miller & Stuble, 2024; Leoschke), we chose to use QR to provide hyperparameters in order to illustrate the automated approach that can be used with such publications do not exist for other species, thus providing a more generally-applicable workflow.

Very few of the studies with spring ephemeral wildflower focus on durations of phenophases, with the notable exception of Miller and Stuble (Miller & Stuble, 2024) who used direct observations of plant phenologies rather that museum records. We investigated several overlapping species: *Erythronium americanum*, *Sanguinaria canadensis*, *Claytonia virginica*, and *Podophyllum peltatum*. Of these shared species, *S. canadensis* and *E. americanum* showed significant decreases in flowering duration in their study; our results show a similar decrease in duration with respect to one or more climate variables.

Lastly, we show the results of the posterior predictive plotting (Fig. 12). The posterior predictive plot provides both a way to summarize the inferred model as well as provide information about the expected patterns in new data. They provide a qualitative way to assess how well the model captures variation in the observations. The example presented in Fig. 12 illustrates how the posterior predictive plot is sensitive to the covariational structure in the covariate data and provides tighter credible intervals when more data are present in the region of covariate space. Detailed summaries of the Bayesian GP results for all 13 species are available in Table S3.

## Discussion

No general statistical theory exists to account for the correspondence between specimen data and latent phenological events. To fill this gap, we derived a unified statistical theory of phenological timing that provides mathematical connections among phenological events: phenological extremes, onsets, phenophase durations, peak phenophase times, cessations, total phenophase time in a population and, most importantly, to the pattern of observed collection times of biological specimens. Based on this theory, we derive a Bayesian inference methodology to recover the hidden states of phenological events from specimen collection times.

Based on this framework, we reached a number of salient conclusions. First, traditional regression methods fail to estimate phenological sensitivities when phenophase durations change systematically with a covariate. As we show, this is because the pattern of specimen collection times is a function of onsets and durations. When the signal from duration patterns is strong and the signal from onset patterns is relatively weak, inferences about timing of onset will be inaccurate. Indeed, traditional regression approaches can recover the wrong sign of a covariate’s effect.

Our methodology addresses this issue. We derived the mathematical function that describes the shape of the distribution of observed collection times (equation [6]). This distribution is not, in general, Gaussian, and is a function of onset times and durations. Our Bayesian methodology finds parameter values (and their uncertainties) that achieve an optimal match with the pattern of observed collection times. Because equation [6] is a function of onset and duration parameters, this inference procedure recovers the values of latent phenological phenomena.

This situation is analogous to earlier comparative studies that did not consider relatedness among species. Correlations between species traits are confounded by the non-independence among species (Felsenstein, 1985). Correlations that were strong before consideration of phylogeny can disappear after the phylogeny is incorporated. When phylogenetic signal is absent, results of conventional regression techniques are valid. Checks like Pagel’s λ (Pagel, 1999) can indicate when phylogenetic non-independence is less of a concern. In parallel, durations confound the estimates of phenological events. When duration is considered in the analysis, patterns in phenological sensitivities can disappear. When duration does not change with respect to the covariates of interest, results of conventional regression techniques are valid. A check for this is analysis of heteroscedasticity; homoscedasticity indicates duration is less likely to be influencing onset model parameter estimates. One consequence of systematic changes in duration is the introduction of heteroscedasticity in the observed-time data. A quick heuristic to assess whether duration may be a confounding factor is to evaluate the degree of heteroscedasticity. If it is substantial, it suggests that variation in duration should be explicitly accounted for to obtain accurate inferences about onset patterns.

Second, we found that duration changed with high posterior probability in relation to at least one covariate for all species examined. This result suggests that the confounding effect of duration is not of marginal significance for species whose phenologies change with covariates.

Third, the methodology opens a new venue of research into phenophase duration based on biocollection data. Phenological mismatch occurs when shifts in phenophases cause interdependent species to fall out of sync. Differences in phenological sensitivity between interacting species can alter the duration and timing of their overlap, potentially leading to trophic mismatch (Edwards & Richardson, 2004; Damien & Tougeron, 2019; Meineke *et al*., 2021), mutualism failure (Warren II & Bradford, 2014), and community restructuring more generally (Miller-Rushing *et al*., 2010). Thus, leveraging museum specimens to study phenophase duration holds significant promise.

Fourth, analyses that fail to include information about population size are likely to provide poor estimates of phenological extremes based on biocollection data, as phenological extremes are functions of population size (equations [8] and [9]). In general, biocollection data do not target phenological events (Davis *et al*., 2015), so the information provided by specimen records is not the same as field observations that target rare or extreme phenological events.

Our methodology addresses this issue by explicitly incorporating information about population size into the analysis. By doing so, the methods recover phenological extreme events with little bias when sample sizes are sufficiently large (e.g., above 60).

Fifth, although there is broad interest in phenophase duration (van Asch & Visser, 2007; Inouye, 2008; Forrest *et al*., 2010; CaraDonna *et al*., 2014; Bock *et al*., 2014; Park *et al*., 2021), very few other methods are available which use biocollection data to estimate duration. Park et al. (Park *et al*., 2025) recently addressed this issue. As far as we can tell, their analysis relied entirely on simulated data, combined with climate samples representative of collection patterns. Despite this, they concluded that herbarium data accurately predict the duration of population-level flowering displays. However, we found that their method confounds estimates of duration and onset: errors in reconstructing these events are correlated, and error magnitude increases with true duration. In contrast, our method yields unbiased duration estimates that are not correlated with errors in onset estimates when priors are unbiased.

Sixth, we developed a theoretical framework to characterize the total range of a phenophase,*RR*, defined as the interval from the first individual to enter the phenophase to the last individual to exit it within a population. We derived the probability density function for this metric, given a specified population size and individual level onset and duration parameters.

Analogous to individual phenophase duration, *RR* captures the window of phenophase activity, but at the population level. Because population sizes are shifting due to anthropogenic pressures, this metric offers a direct means to assess how such changes affect the temporal extent of phenophase activity in a population. We therefore propose that *RR* is a key metric for addressing phenological mismatch in the context of anthropogenic change.

Seventh, this methodology provides a way to investigate variability in phenological processes by decomposing it into components attributable to onset and to duration. Although we did not explore inferences about variability in depth, we suggest that such considerations can lead to phenological paradoxes similar to those observed in onset inference. In particular, the width of the observed-time distribution, *T*, is not a direct proxy for phenophase duration. While the observed-time range must be at least as wide as the longest individual duration in the population, variability in onset times also contributes to its spread. Thus, the observed-time distribution reflects a confounding interplay between onset and duration variability. This concept is illustrated in Fig. 13.

### Future Directions

We introduce a broad, unified statistical framework for modeling phenological distributions. However, several important issues remain beyond the scope of this study, including the treatment of biases, outliers, and the role of σ. The Bayesian GP methodology relies on several statistical assumptions that may be violated in practice. Many of these are similar to assumptions underlying parametric statistical tests, but one in particular deserves further attention: the assumption that collection events represent a random sample of times within a phenophase. In reality, it is well-known that herbarium and other natural history collections are often biased due to the practices and preferences of taxonomists and collectors (Iler *et al*., 2017; Daru *et al*., 2018; Jones & Daehler, 2018; Panchen *et al*., 2019; Pearson, 2019c; Taylor, 2019; Belitz *et al*., 2020; Zizka *et al*., 2021; Park *et al*., 2023; Lai, 2025). We don’t believe that collection biases will substantially impact inferences using Bayesian GP any more than they would impact inferences using most other statistical methodologies like linear regression. Park et al. (Park *et al*., 2025) used simulations to address questions of collection bias and accuracy of their method. This pioneering study presents a workflow to likewise investigate the influence of biases using Bayesian GP. (Moreover, we advocate using their quantile regression approach to initialize prior hyperparameters for Bayesian GP.)

A second issue concerns model complexity. We used standard linear models to describe the relationship between covariates and phenological parameters; however, these may not adequately capture more complex patterns in the data. While we examined how model mismatch affects inferences in the absence of covariates, we did not investigate inference errors that may arise when covariate effects deviate from linear assumptions. This is another avenue of research worth exploring.

We hope that this theory and methodology will encourage broader application of museum specimens to address fundamental questions in ecology and evolutionary biology.

## Acknowledgements

Thank you to Anne Estes, Lucinda McDade, and Katy Adams for providing reviews of earlier versions of the manuscript and for providing additional insights. Special thanks to Martin Modrák for theoretical insights, for providing the GP approach, and for providing a C++ function that compiles with Stan code and increases computational efficiency dramatically over prior incarnations.

## Author Contribution

Conceptualization: DJH, DC; R programming and packaging: DJH, DC; Stan programming: DJH; Mathematical derivations: DJH; Writing the manuscript: DJH; Processing empirical datasets: DJH.

## Supporting Information

Additional Supporting Information may be found online in the Supporting Information section at the end of the article.

Table S1. Simulation parameters.

Table S2. Summary statistics for the empirical dataset

Table S3. Results of the Bayesian GP analysis

## Notes

### Competing Interest Statement

The authors have declared no competing interest.

## References

1. Abu-Asab MS, Peterson PM, Shetler SG, Orli SS. 2001. Earlier plant flowering in spring as a response to global warming in the Washington, DC, area. Biodiversity & Conservation 10: 597–612.

2. Aggarwal CC. 2017. Outlier Analysis. Cham: Springer.

3. Ahlstrand NI, Primack RB, Austin MW, Panchen ZA, Römermann C, Miller-Rushing AJ. 2025. The promise of digital herbarium specimens in large-scale phenology research. New Phytologist: nph.70178.

4. Alecrim EF, Sargent RD, Forrest JRK. 2023. Higher-latitude spring-flowering herbs advance their phenology more than trees with warming temperatures. Journal of Ecology 111: 156–169.

5. Amano T, Smithers RJ, Sparks TH, Sutherland WJ. 2010. A 250-year index of first flowering dates and its response to temperature changes. Proceedings of the Royal Society B: Biological Sciences 277: 2451– 2457.

6. van Asch M, Visser ME. 2007. Phenology of forest caterpillars and their host trees: the importance of synchrony. Annual Review of Entomology 52: 37–55.

7. Badeck F, Bondeau A, Böttcher K, Doktor D, Lucht W, Schaber J, Sitch S. 2004. Responses of spring phenology to climate change. New Phytologist 162: 295–309.

8. Bates JM, Fidino M, Nowak-Boyd L, Strausberger BM, Schmidt KA, Whelan CJ. 2023. Climate change affects bird nesting phenology: Comparing contemporary field and historical museum nesting records. Journal of Animal Ecology 92: 263–272.

9. Beauvais M-P, Pellerin S, Dubé J, Lavoie C. 2017. Herbarium specimens as tools to assess the impact of large herbivores on plant species. Botany 95: 153–162.

10. Belitz MW, Larsen EA, Ries L, Guralnick RP. 2020. The accuracy of phenology estimators for use with sparsely sampled presence-only observations (A Ellison, Ed.). Methods in Ecology and Evolution 11: 1273–1285.

11. Betancourt M. 2017. A Conceptual Introduction to Hamiltonian Monte Carlo. arXiv.org. 10.48550/arXiv.1701.02434

12. Betancourt MJ, Girolami M. 2013. Hamiltonian Monte Carlo for Hierarchical Models. DOI:10.1201/b18502-5

13. Bishop CM. 2006. Pattern Recognition and Machine Learning. New York: Springer.

14. Blair JM, Edwards CA, Johnson JH. 1976. Rational Chebyshev Approximations for the Inverse of the Error Function. Mathematics of Computation 30: 827–862.

15. Bock A, Sparks TH, Estrella N, Jee N, Casebow A, Schunk C, Leuchner M, Menzel A. 2014. Changes in first flowering dates and flowering duration of 232 plant species on the island of Guernsey. Global Change Biology 20: 3508–3519.

16. Büntgen U, Piermattei A, Krusic PJ, Esper J, Sparks T, Crivellaro A. 2022. Plants in the UK flower a month earlier under recent warming. Proceedings of the Royal Society B: Biological Sciences 289: 20212456.

17. Calinger KM, Queenborough S, Curtis PS. 2013. Herbarium specimens reveal the footprint of climate change on flowering trends across north-central North America (Y Buckley, Ed.). Ecology Letters 16: 1037–1044.

18. CaraDonna PJ, Iler AM, Inouye DW. 2014. Shifts in flowering phenology reshape a subalpine plant community. Proceedings of the National Academy of Sciences of the United States of America 111: 4916–4921.

19. Carpenter B, Gelman A, Hoffman MD, Lee D, Goodrich B, Betancourt M, Brubaker M, Guo J, Li P, Riddell A. 2017. *Stan*: A Probabilistic Programming Language. Journal of Statistical Software 76.

20. Cody WJ. 1993. Algorithm 715: SPECFUN–a portable FORTRAN package of special function routines and test drivers. ACM Transactions on Mathematical Software 19: 22–30.

21. Cook RD. 1977. Detection of Influential Observation in Linear Regression. Technometrics 19: 15-18.

22. Cook RD, Weisberg S. 1995. Residuals and influence in regression. New York: Chapman & Hall.

23. Cooke P. 1979. Statistical inference for bounds of random variables. Biometrika 66: 367–374.

24. Cooke P. 1980. Optimal linear estimation of bounds of random variables. Biometrika 67: 257–258.

25. Crimmins TM. 2021. The USA National Phenology Network: Big Idea, Productivity, and Potential—and Now, at Big Risk. The Bulletin of the Ecological Society of America 102: e01802.

26. Damien M, Tougeron K. 2019. Prey–predator phenological mismatch under climate change. Current Opinion in Insect Science 35: 60–68.

27. Daru BH, Park DS, Primack RB, Willis CG, Barrington DS, Whitfeld TJS, Seidler TG, Sweeney PW, Foster DR, Ellison AM, et al. 2018. Widespread sampling biases in herbaria revealed from large-scale digitization. New Phytologist 217: 939–955.

28. Davis CC, Willis CG, Connolly B, Kelly C, Ellison AM. 2015. Herbarium records are reliable sources of phenological change driven by climate and provide novel insights into species’ phenological cueing mechanisms. American Journal of Botany 102: 1599–1609.

29. Edery I. 2000. Circadian rhythms in a nutshell. Physiological Genomics 3: 59–74.

30. Edwards W, Lindman H, Savage LJ. 1963. Bayesian statistical inference for psychological research. Psychological Review 70: 193–242.

31. Edwards M, Richardson AJ. 2004. Impact of climate change on marine pelagic phenology and trophic mismatch. Nature 430: 881–884.

32. Elfving G. 1947. THE ASYMPTOTICAL DISTRIBUTION OF RANGE IN SAMPLES FROM A NORMAL POPULATION. Biometrika 34: 111–119.

33. Faidiga AS, Oliver MG, Budke JM, Kalisz S. 2023. Shifts in flowering phenology in response to spring temperatures in eastern Tennessee. American Journal of Botany 110: e16203.

34. Felsenstein J. 1985. Phylogenies and the Comparative Method. The American Naturalist 125: 1–15.

35. Fitchett JM, Grab SW, Thompson DI. 2015. Plant phenology and climate change: Progress in methodological approaches and application. Progress in Physical Geography: Earth and Environment 39: 460–482.

36. Fitter AH, Fitter RSR. 2002. Rapid Changes in Flowering Time in British Plants. Science 296: 1689–1691.

37. Forrest J, Inouye DW, Thomson JD. 2010. Flowering phenology in subalpine meadows: does climate variation influence community co-flowering patterns? Ecology 91: 431–440.

38. Gabry J, Češnovar R, Johnson A, Bronder S. 2025. cmdstanr: R Interface to ‘CmdStan’.

39. Hastings WK. 1970. Monte Carlo Sampling Methods Using Markov Chains and Their Applications. Biometrika 57: 97–109.

40. Hearn D. 2022. A general theory of phenological timing reveals counterintuitive interpretations of historical patterns of phenological change.

41. Heberling JM. 2022. Herbaria as Big Data Sources of Plant Traits. International Journal of Plant Sciences 183: 87–118.

42. Helm B, Ben-Shlomo R, Sheriff MJ, Hut RA, Foster R, Barnes BM, Dominoni D. 2013. Annual rhythms that underlie phenology: biological time-keeping meets environmental change. Proceedings of the Royal Society B: Biological Sciences 280: 20130016.

43. Hofert M, Kojadinovic I, Maechler M, Yan J. 2025. copula: Multivariate Dependence with Copulas.

44. Hollister J, Shah T, Nowosad J, Robitaille A, Beck M, Johnson M. 2023. elevatr: Access Elevation Data from Various APIs.

45. Iler AM, Humphrey PT, Ogilvie JE, CaraDonna PJ. 2021. Conceptual and practical issues limit the utility of statistical estimators of phenological events. Ecosphere 12: e03828.

46. Iler AM, Inouye DW, Schmidt NM, Høye TT. 2017. Detrending phenological time series improves climate-phenology analyses and reveals evidence of plasticity. Ecology 98: 647–655.

47. Inouye DW. 2008. Effects of climate change on phenology, frost damage, and floral abundance of montane wildflowers. Ecology 89: 353–362.

48. Iwanycki Ahlstrand N, Primack RB, Tøttrup AP. 2022. A comparison of herbarium and citizen science phenology datasets for detecting response of flowering time to climate change in Denmark. International Journal of Biometeorology 66: 849–862.

49. Jaynes ET. 2021. Probability Theory: The Logic of Science (GL Bretthorst, Ed.). Cambridge: Cambridge University Press.

50. Jones CA, Daehler CC. 2018. Herbarium specimens can reveal impacts of climate change on plant phenology; a review of methods and applications. PeerJ 6: e4576.

51. Lai HR. 2025. Model-based ordination for phenological studies: From controlling sampling bias to inferring temporal associations. Methods in Ecology and Evolution: 2041–210X.70079.

52. Lavoie C. 2013. Biological collections in an ever changing world: Herbaria as tools for biogeographical and environmental studies. *Perspectives in Plant Ecology*, Evolution and Systematics 15: 68–76.

53. Lavoie C, Lachance D. 2006. A new herbarium-based method for reconstructing the phenology of plant species across large areas. American Journal of Botany 93: 512–516.

54. Leoschke MJ. Dodecatheon meadia, in Iowa tallgrass prairie.

55. Li X, Deng S, Li L, Jiang Y. 2019. Outlier Detection Based on Robust Mahalanobis Distance and Its Application. Open Journal of Statistics 9: 15–26.

56. Mazer SJ, Love NLR, Park IW, Ramirez-Parada T, Matthews ER. 2021. PHENOLOGICAL SENSITIVITIES TO CLIMATE ARE SIMILAR IN TWO CLARKIA CONGENERS: INDIRECT EVIDENCE FOR FACILITATION, CONVERGENCE, NICHE CONSERVATISM, OR GENETIC CONSTRAINTS. Madroño 68: 388–405.

57. McElreath R. 2020. Statistical Rethinking: A Bayesian Course with Examples in R and STAN. CRC Press.

58. Meineke EK, Davies TJ. 2019. Museum specimens provide novel insights into changing plant–herbivore interactions. Philosophical Transactions of the Royal Society B: Biological Sciences 374: 20170393.

59. Meineke EK, Davis CC, Davies TJ. 2021. Phenological sensitivity to temperature mediates herbivory. Global Change Biology 27: 2315–2327.

60. Metropolis N, Rosenbluth AW, Rosenbluth MN, Teller AH, Teller E. 1953. Equation of State Calculations by Fast Computing Machines. The Journal of Chemical Physics 21: 1087–1092.

61. Michonneau F, Collins M, Chamberlain S. 2016. ridigbio: An interface to iDigBio’s search API that allows downloading specimen records.

62. Miller TK, Heberling JM, Kuebbing SE, Primack RB. 2023. Warmer temperatures are linked to widespread phenological mismatch among native and non-native forest plants. Journal of Ecology 111: 356–371.

63. Miller CN, Stuble KL. 2024. Warm Spring Days are Related to Shorter Durations of Reproductive Phenophases for Understory Forest Herbs. Ecology and Evolution 14: e70700.

64. Miller-Rushing AJ, Høye TT, Inouye DW, Post E. 2010. The effects of phenological mismatches on demography. Philosophical Transactions of the Royal Society B: Biological Sciences 365: 3177–3186.

65. Miller-Rushing AJ, Lloyd-Evans TL, Primack RB, Satzinger P. 2008. Bird migration times, climate change, and changing population sizes. Global Change Biology 14: 1959–1972.

66. Miller-Rushing AJ, Primack RB, Primack D, Mukunda S. 2006. Photographs and herbarium specimens as tools to document phenological changes in response to global warming. American Journal of Botany 93: 1667–1674.

67. Mood, A.M. 1974. Introduction to the theory of statistics. McGraw-Hill Education – Europe.

68. Moussus J, Julliard R, Jiguet F. 2010. Featuring 10 phenological estimators using simulated data. Methods in Ecology and Evolution 1: 140–150.

69. Neal RM. 2011. MCMC using Hamiltonian dynamics.

70. Oldham MJ. 1990. STATUS REPORT ON THE WILD HYACINTH CAMASSIA SCILLOIDES.

71. Pagel M. 1999. Inferring the historical patterns of biological evolution. Nature 401: 877–884.

72. Panchen ZA, Doubt J, Kharouba HM, Johnston MO. 2019. Patterns and biases in an Arctic herbarium specimen collection: Implications for phenological research. Applications in Plant Sciences 7: e01229.

73. Panchen ZA, Primack RB, Aniśko T, Lyons RE. 2012. Herbarium specimens, photographs, and field observations show Philadelphia area plants are responding to climate change. American Journal of Botany 99: 751–756.

74. Park DS, Breckheimer I, Williams AC, Law E, Ellison AM, Davis CC. 2019. Herbarium specimens reveal substantial and unexpected variation in phenological sensitivity across the eastern United States. Philosophical Transactions of the Royal Society B: Biological Sciences 374: 20170394.

75. Park DS, Lyra GM, Ellison AM, Maruyama RKB, Dos Reis Torquato D, Asprino RC, Cook BI, Davis CC. 2023. Herbarium records provide reliable phenology estimates in the understudied tropics. Journal of Ecology 111: 327–337.

76. Park IW, Mazer SJ. 2018. Overlooked climate parameters best predict flowering onset: Assessing phenological models using the elastic net. Global Change Biology 24: 5972–5984.

77. Park IW, Ramirez-Parada T, Mazer SJ. 2021. Advancing frost dates have reduced frost risk among most North American angiosperms since 1980. Global Change Biology 27: 165–176.

78. Park IW, Ramirez-Parada T, Record S, Davis C, Ellison AM, Mazer SJ. 2025. Herbarium data accurately predict the timing and duration of population-level flowering displays. Ecography 2025: e06961.

79. Pearse WD, Davis CC, Inouye DW, Primack RB, Davies TJ. 2017. A statistical estimator for determining the limits of contemporary and historic phenology. Nature Ecology & Evolution 1: 1876–1882.

80. Pearson KD. 2019a. A new method and insights for estimating phenological events from herbarium specimens. Applications in Plant Sciences 7: e01224.

81. Pearson KD. 2019b. A new method and insights for estimating phenological events from herbarium specimens. Applications in Plant Sciences 7.

82. Pearson KD. 2019c. Spring– and fall-flowering species show diverging phenological responses to climate in the Southeast USA. International Journal of Biometeorology 63: 481–492.

83. Petrauski L, Owen SF, Constantz GD, Anderson JT. 2019. Changes in flowering phenology of Cardamine concatenata and Erythronium americanum over 111 years in the Central Appalachians. Plant Ecology 220: 817–828.

84. Pierce D. 2025. Interface to Unidata netCDF (Version 4 or Earlier) Format Data.

85. Primack RB, Gallinat AS, Ellwood ER, Crimmins TM, Schwartz MD, Staudinger MD, Miller-Rushing AJ. 2023. Ten best practices for effective phenological research. International Journal of Biometeorology 67: 1509–1522.

86. Primack D, Imbres C, Primack RB, Miller-Rushing AJ, Del Tredici P. 2004a. Herbarium specimens demonstrate earlier flowering times in response to warming in Boston. American Journal of Botany 91: 1260–1264.

87. Primack D, Imbres C, Primack RB, Miller-Rushing AJ, Del Tredici P. 2004b. Herbarium specimens demonstrate earlier flowering times in response to warming in Boston. American Journal of Botany 91: 1260–1264.

88. R Core Team. 2025. R: A Language and Environment for Statistical Computing.

89. Raible F, Takekata H, Tessmar-Raible K. 2017. An Overview of Monthly Rhythms and Clocks. Frontiers in Neurology 8: 189.

90. Ramirez-Parada TH, Park IW, Record S, Davis CC, Ellison AM, Mazer SJ. 2024. Plasticity and not adaptation is the primary source of temperature-mediated variation in flowering phenology in North America. Nature Ecology & Evolution 8: 467–476.

91. Rank J. 2007. Copulas: From theory to application in Finance. London: Bloomberg Press.

92. Rasmussen CE, Williams CKI. 2008. Gaussian Processes for Machine Learning. Cambridge, Mass.: The MIT Press.

93. Renner SS, Wesche M, Zohner CM. 2021. Climate data and flowering times for 450 species from 1844 deepen the record of phenological change in southern Germany. American Journal of Botany 108: 711– 717.

94. Renner SS, Zohner CM. 2018. Climate Change and Phenological Mismatch in Trophic Interactions Among Plants, Insects, and Vertebrates. Annual Review of Ecology, Evolution, and Systematics 49: 165–182.

95. Robbirt KM, Davy AJ, Hutchings MJ, Roberts DL. 2011. Validation of biological collections as a source of phenological data for use in climate change studies: a case study with the orchid Ophrys sphegodes: Herbarium specimens for climate change studies. Journal of Ecology 99: 235–241.

96. Rohde RA, Hausfather Z. 2020. The Berkeley Earth Land/Ocean Temperature Record. Earth System Science Data 12: 3469–3479.

97. Sherry RA, Zhou X, Gu S, Arnone JA, Schimel DS, Verburg PS, Wallace LL, Luo Y. 2007. Divergence of reproductive phenology under climate warming. Proceedings of the National Academy of Sciences 104: 198–202.

98. Taylor SD. 2019. Estimating flowering transition dates from status-based phenological observations: a test of methods. PeerJ 7: e7720.

99. Thackeray SJ, Henrys PA, Hemming D, Bell JR, Botham MS, Burthe S, Helaouet P, Johns DG, Jones ID, Leech DI, et al. 2016a. Phenological sensitivity to climate across taxa and trophic levels. Nature 535: 241–245.

100. Thackeray SJ, Henrys PA, Hemming D, Bell JR, Botham MS, Burthe S, Helaouet P, Johns DG, Jones ID, Leech DI, et al. 2016b. Phenological sensitivity to climate across taxa and trophic levels. Nature 535: 241–245.

101. Wang H, Ge Q, Rutishauser T, Dai Y, Dai J. 2015. Parameterization of temperature sensitivity of spring phenology and its application in explaining diverse phenological responses to temperature change. Scientific Reports 5: 8833.

102. Warren II RJ, Bradford MA. 2014. Mutualism fails when climate response differs between interacting species. Global Change Biology 20: 466–474.

103. Watson MA, Vuorisalo T. 2024. Interactions between developmental phenology, carbon movement, and storage constrain demography in the understory clonal herb Podophyllum peltatum L. Frontiers in Plant Science 15: 1325052.

104. Willems FM, Scheepens JF, Bossdorf O. 2022. Forest wildflowers bloom earlier as Europe warms: lessons from herbaria and spatial modelling. New Phytologist 235: 52–65.

105. Williams DA. 1987. Generalized Linear Model Diagnostics Using the Deviance and Single Case Deletions. Applied Statistics 36: 181.

106. Williams TM, Schlichting CD, Holsinger KE. 2021. Herbarium records demonstrate changes in flowering phenology associated with climate change over the past century within the Cape Floristic Region, South Africa. Climate Change Ecology 1: 100006.

107. Willis CG, Ellwood ER, Primack RB, Davis CC, Pearson KD, Gallinat AS, Yost JM, Nelson G, Mazer SJ, Rossington NL, et al. 2017. Old Plants, New Tricks: Phenological Research Using Herbarium Specimens. Trends in Ecology & Evolution 32: 531–546.

108. Wilson SM, Anderson JH, Ward EJ. 2023. Estimating phenology and phenological shifts with hierarchical modeling. Ecology 104: e4061.

109. Xie Y, Thammavong HT, Park DS. 2022. The ecological implications of intra– and inter-species variation in phenological sensitivity. New Phytologist 236: 760–773.

110. Zizka A, Antonelli A, Silvestro D. 2021. *sampbias*, a method for quantifying geographic sampling biases in species distribution data. Ecography 44: 25–32.

111. Zohner CM, Mo L, Renner SS. 2018. Global warming reduces leaf-out and flowering synchrony among individuals. eLife 7: e40214.

